# Morphogenetic constrains in the development of gastruloids: implications for mouse gastrulation

**DOI:** 10.1101/2024.12.12.628151

**Authors:** U.M. Fiuza, S. Bonavia, P. Pascual-Mas, G. Torregrosa-Cortés, P. Casaní-Galdón, G. Robertson, A. Dias, A. Martinez Arias

## Abstract

Mammalian embryonic size is tightly controlled with checkpoints and compensatory mechanisms correcting size defects. Here, we take advantage of gastruloids, a stem cell embryoid system not subject to most size controls, to study the role of size in emergent properties of mammalian embryogenesis. We report that gastruloids exhibit robust morphology and transcriptional profiles within a size range. However, size affects the dynamics, and, outside a range of robust morphogenesis, the precision of anterior-posterior (AP) axial elongation. Gastruloid axial elongation exhibits active cellular contractility, requires planar cell polarity (PCP), adhesion and cell-cell contact remodelling. Smaller gastruloids initiate elongation earlier, correlated with an earlier Brachyury polarisation. Brachyury expression increases tissue fluidity. Axis formation is regulated by the balance of Brachyury multifoci coalescence and the timing of initiation of the elongation programme. Sizes beyond the robust range can modify relative tissue composition. Very small aggregates have increased neural fate bias, accompanied by a loss of paraxial mesoderm mediated by differences in Nodal signalling activity.

## Introduction

A capacity to keep a proportionate body plan across a range of sizes is an important feature of animal embryos. In 1892 Hans Driesch separated the first two blastomeres of a sea urchin embryo and let them develop (Driesch, 1892). To his surprise, he obtained two fully fledged and well proportioned larvae but each about half the size than the one that would have emerged if the cells had been left together (Driesch, 1892, Driesch, 1902). A comparable experiment was repeated in frogs with a similar result (McClendon, 1910). Mammals exhibit a similar behaviour during blastocyst formation (Sainz et al., 2016) but have compensatory mechanisms afterwards (Snow and Tam, 1979; Power and Tam, 1993; Saiz et al, 2016; Martinez Arias et al 2013; Lewis and Rossant, 1982).

The mechanisms and consequences associated with the development of different sized embryos are difficult to study, particularly in mammalian embryos, where development takes places in the uterus. However, over the last few years the development of pluripotent stem cell (PSC) models of early embryogenesis (Shahbazi et al., 2019; Turner and Martinez Arias, 2024) has opened up experimental possibilities that can be used to explore aspects of this question. One of these models, the gastruloid system, is proving experimentally versatile. Gastruloids are aggregates of PSCs that upon defined culture conditions develop the main features of the mammalian body plan in a manner, and time course, that mimics the process of gastrulation. Specifically, upon a short pulse of the Wnt signalling agonist Chiron (CHIR) they self-organise into a polarised structure that elongates along an anteroposterior axis with patterns and sequences of gene expression that mimic those of the embryo during and after gastrulation (Van der Brink et al., 2014; Beccari et al., 2018). Gastruloids are robust and reproducible (Turner et al., 2017; Merle et al., 2024), and this is the reason why they are being used, as a complement to embryos, to probe in detail mechanisms of symmetry breaking, the control of gene expression or features of cell collectives during early embryogenesis (Moris et al, 2020; Turner and Martinez Arias, 2024).

The initial number of cells in the culture is a crucial variable for the development of gastruloids. ‘Chiron driven gastruloids’ form properly within a range of cell numbers (between about 40 and 300), over and above which abnormalities ensue (van den Brink et al, 2014). They exhibit very precise spatial patterns of gene expression organisation across the working size range that, most surprisingly, scale with great precision (Merle et al, 2024). Here we have exploited this property of gastruloids to study the influence of size on the organisation of gene expression and morphogenesis.

Our results confirm that, within a size range, the cell type composition in the gastruloids is stable, but define precise boundaries to these numbers and observe that the dynamics of the emergence of an antero-posterior axis varies, with smaller gastruloids initiating this process earlier than larger ones. We correlate this observation with the onset of *TbxT/Brachyury* (*TbxT*) polarisation, whose expression becomes first restricted to discrete domains that over time coalesce into one. Outside the working size range, in larger gastruloids the multiple axes that appear correlate with the difficulty of multiple foci of TbxT expression to coalesce into a single domain. We observe that these TbxT foci are enriched in E-cadherin (E-Cdh) and suggest that sorting through differential adhesion could be a driving mechanism of such coalescence in a manner that relates to size. A similar sorting behaviour has been reported in the establishment of Wnt signalling domains in gastruloids (McNamara et al, 2024) and both might be related. We also observe that the smallest gastruloids, besides failing to elongate reliably, exhibit a bias towards neural fates that correlates with the levels of Nodal expression.

Our results show a significant effect of size on the spatial organisation of gene expression during embryogenesis and suggest that there might be mechanisms that coordinate signalling, transcription and morphogenesis to some global references associated with size.

## Results

### Robustness of gastruloid development

The development of gastruloids relies on a number of variables most important of which are the Pluripotent Stem Cell (PSC) line and the culture conditions before aggregation (Turner and Martinez Arias, 2024) (Figure 1A and S2). In our experiments, we use E14 cells grown in Serum and LIF. Under these conditions and in agreement with previous observations, after the pulse of Chiron, gastruloids exhibit a round shape at 72h (Figure S1A), ellipsoid at 96h (Figure S1B) and are elongated by 120h (Figure 1A). Seeding different initial numbers of cells, gastruloids form mono axially elongating structures robustly within a limited size range, and we were able to define this range precisely between 75 and 300 cells (Figures 1A-D). The size range, when defined by the initial cell seeding number, is cell line dependent (Figure S2A). Gastruloids below a size threshold will exhibit variable elongation, with increasing sizes above 300 starting cells exhibit increasing numbers of structures with multi axes (Figure 1B-D).

**Figure 1.**
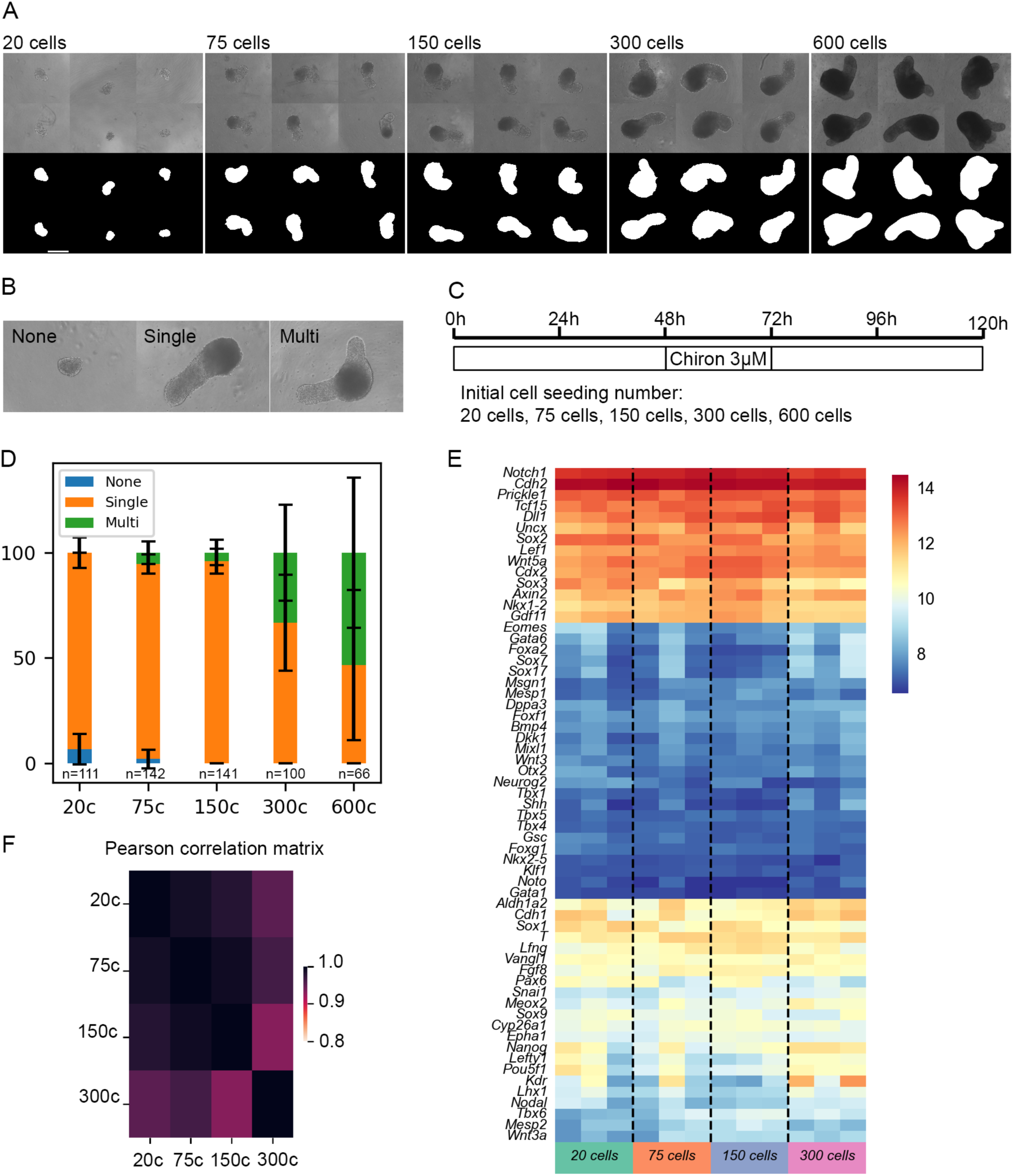
Gastruloids are transcriptionally and morphologically robust over a size range. (A) Morphological features of gastruloids ranging from 20 cells to 600 cells (initial cell seeding number) at 120h. (B) Gastruloid axis formation morphological classes at 120h. (C) Scheme of experimental protocol for producing gastruloids. (D) Quantification of gastruloid axis formation phenotypes across sizes at 120h. N=3 to 4 replicates. (E) Heatmap of expression profiles of developmental genes across gastruloids of different sizes at 120h (from bulk mRNAseq). N=3 replicates. (F) Pearson correlation score matrix comparing the expression of fate markers and other developmental genes between gastruloids of different sizes at 120h (from bulk mRNAseq; Table S1). N=3 replicates. 20c - 20 cells, 75c - 75 cells, 150c – 150 cells, 300c – 300 cells (initial cell seeding number; this notation is repeated in subsequent figures).

We performed bulk mRNA sequencing analysis of gastruloids of different sizes and observe, they exhibit overall transcriptional robustness in developmental genes (Figure 1E-F). This observation is consistent with a previous report on the precision and scalability of gene expression in gastruloids (Merle et al, 2024). At the whole transcriptome level and across genes associated with different specific cell types, the transcriptional expression profiles are globally well conserved (Figure 1F-E). However, in a comparison of DGE (differential gene expression analysis) between extreme gastruloid sizes (20-cell vs 300-cell gastruloids) we observe differences in the expression of some cell type specific genes (see also Figure 6A, discussed below).

We wondered if this robustness of gene expression to size corresponded to a similar robustness in the first morphogenetic event associated with gastruloid development: elongation.

### Gastruloids require active cytoskeleton activity and Wnt-mediated PCP for AP axial elongation

In a comparison of the timing of elongation in gastruloids with different sizes, we observe that smaller gastruloids initiate elongation earlier than larger gastruloids (Figure 2A, movie 1). This behaviour is conserved across cell lines with different genetic backgrounds (Figure S3A and S3B) and correlates with the timing of the polarisation of *TbxT/Bra* expression (Figure 2C-D). Furthermore, *TbxT* is required for elongation as gastruloids from *TbxT* mutant cells do not elongate (Figure 2B; Wehmeyer et al., 2022).

**Figure 2.**
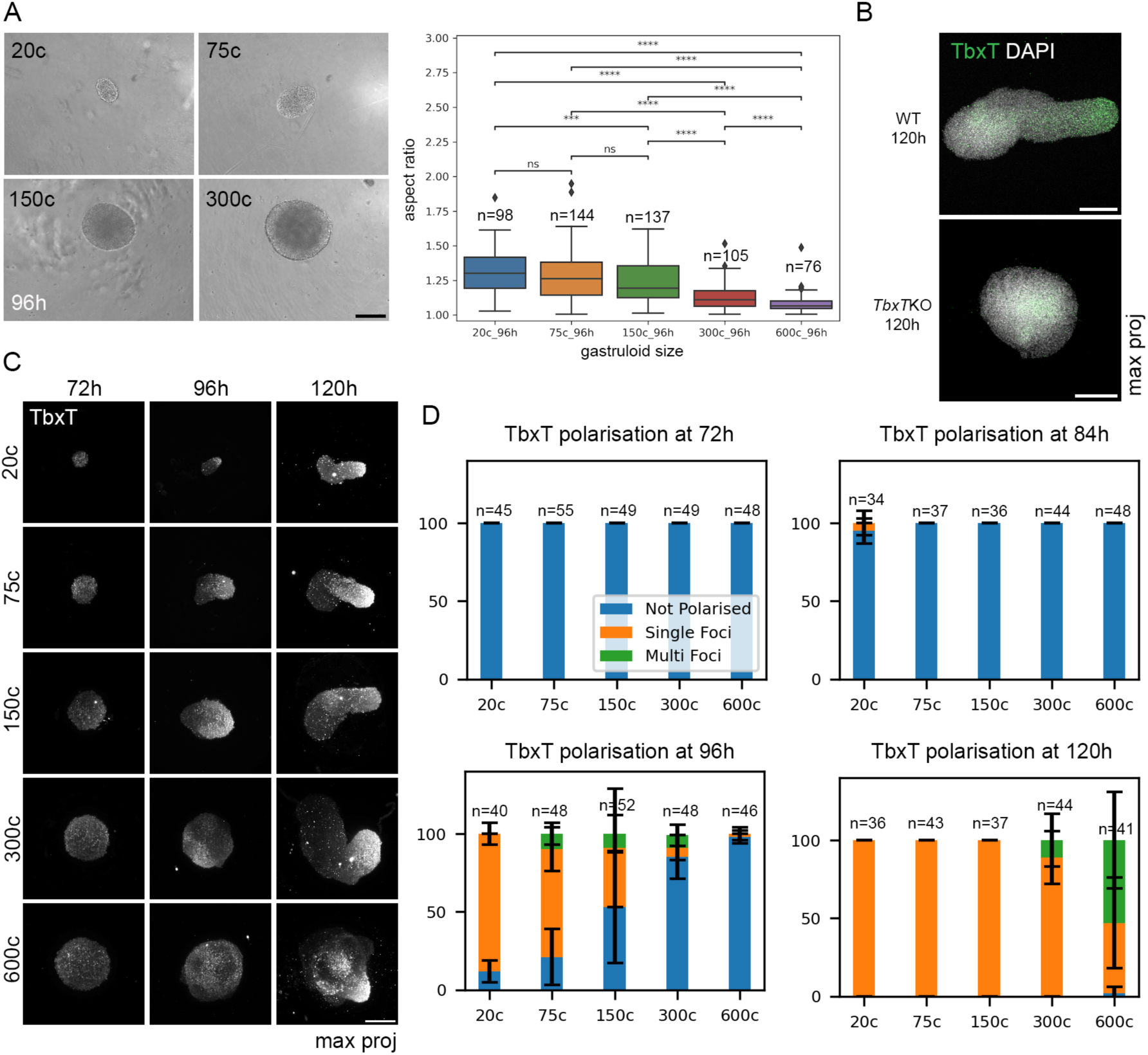
Smaller gastruloids initiate AP elongation earlier which correlates with an earlier polarisation of TbxT. (A) Morphologies of gastruloids with different sizes at 96h. Quantification of the aspect ratio of gastruloids with different sizes at 96h. N=3 to 4 replicates. (B) HCR (hairpin chain reaction) labelling of TbxT in 120h WT and TbxT^-/-^ (TbxTKO) gastruloids. (C) Immunofluorescences characterising the evolution of the pattern of TbxT expression in gastruloids of different sizes at 72h, 96h and 120h. (D) Quantification of the degree of polarization of the TbxT expression, i.e., heterogeneous expression – not polarised, single foci – polarised in a single domain, and multi foci – polarised in more than a domain. N=3 replicates. Scale bar – 200μm.

It has been suggested that the elongation of gastruloids is related to a process of Convergent Extension (CE) (de Jong et al, 2024). To test this, we analysed cell motion in gastruloids with sparse-labelled nuclei using light sheet microscopy and non-linear registration (see Material and Methods). The nuclear movement analysis was performed using a custom-made pipeline based on image registration (for more details see Material and Methods and Figure S20). In this experiment, we used three different metrics to characterise cell movement: cumulative path, drift (directionality modulus) and diffusion (mobility metric) (Figure 3A, Figure S2B, movie2 and movie3). Cells located in the elongating region of the gastruloid exhibit greater motility than those located at the opposite, anterior, one. This is evident from the higher cumulative path and diffusion of these cells. There is as well as some increased directionality, reflected in their higher drift. Cells extruded by the gastruloid can be seen outside the sample with perceived high motility (Figure 3A).

**Figure 3.**
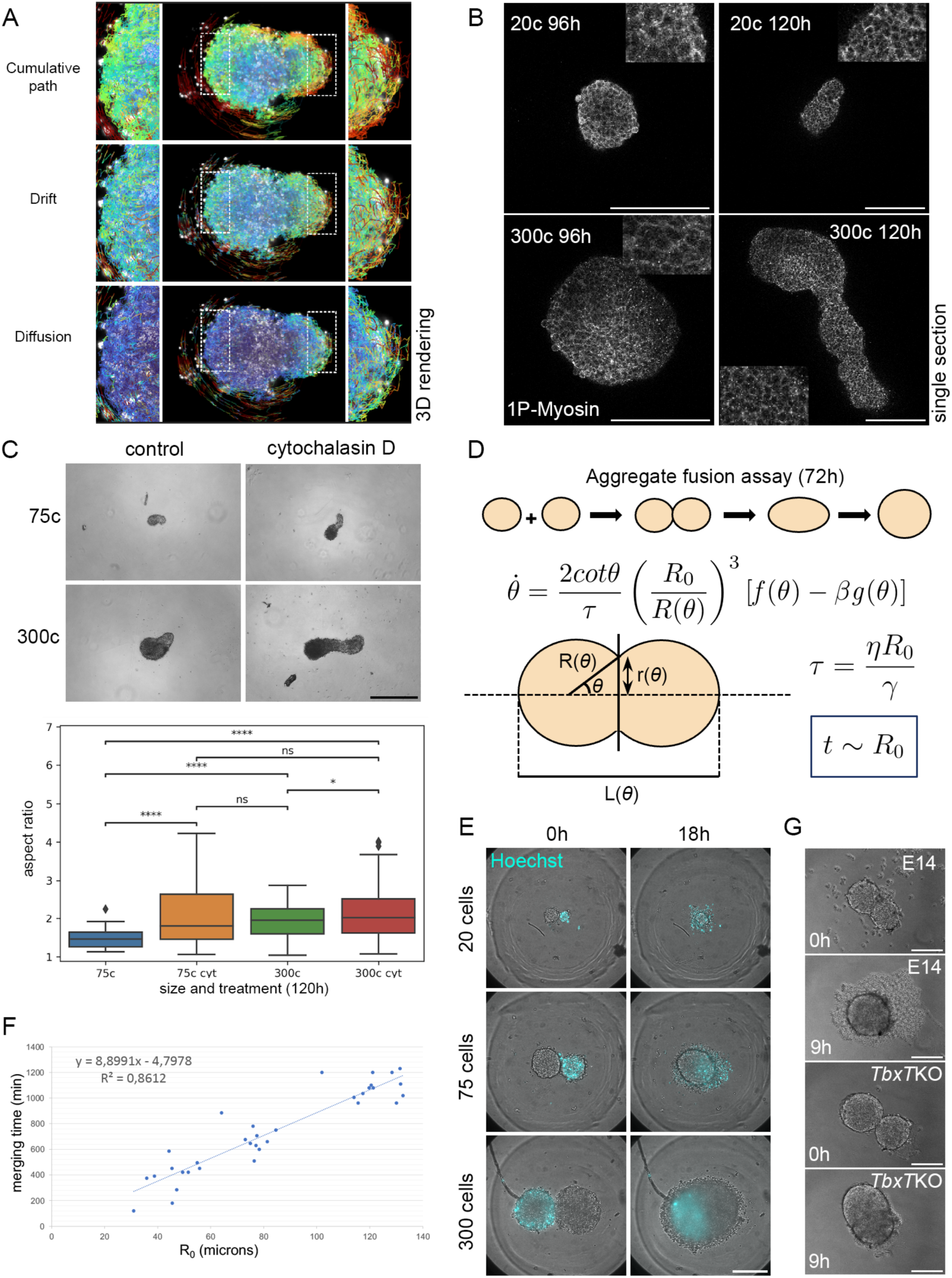
Mechanics of gastruloid elongation. (A) Characterisation of tissue cellular movement dynamics in a 75-cell gastruloid with light sheet microscopy and image analysis (non-linear registration regional cell movement analysis). Data derived from a chimeric gastruloid with nuclear sparse-labelling (60% E14 cells + 40% emiRFP670 cells). Posterior cells present the highest level of movement in the gastruloid. The high movement of extruded cells results in part of an artifact that as these cells become attached to the imaging chamber, rigid registration corrections for the gastruloid movement will artificially generate a higher perception of relative movement in the extruded cells uncoupled to the gastruloid. Anterior to the left and posterior to the right. Red – max values; blue - min values. (B) PhosphoMyosin Light Chain 2, Ser19 (marker of actively contracting myosin) immunostaining of gastruloids of 20 cells and 300 cells at 96h and 120h. (C) Effects of a 0,05ng/ml treatment of cytochalasin D during the elongation period (90h-120h) on gastruloid elongation at 120h. Brightfield images and quantification of aspect ratio in control and treated gastruloids (sizes: 75 cells and 300 cells). N = 3 replicates. (D) Force balance mechanical model of aggregate fusion adapted from Oriola et al, 2020. Inference made from the model: if the tissue mechanical properties are the same across different sizes, then the aggregate merging time should be directly proportional to the initial radius of the aggregates. *θ* - fusion angle, *R*(*θ*) - radius of the aggregates, *r*(*θ*) - neck radius during fusion, *L*(*θ*) - total length of the two aggregates, *β* - elastocapillary number, i.e., dimensionless parameter quantifying the degree of fusion, *τ* - viscocapillary time. (E) Merging assays with aggregates of different sizes where one aggregate is labelled with Hoechst and the other is unlabelled, at 0h and 18h after being put together. Merging experiments taking place in a Viventis LS1 imaging chamber. N = 3 replicates. (F) Analysis of a linear relationship between the initial aggregate radius and the time required for two aggregates to merge (defined as the time at which the merging aggregates present the highest circularity). N = 3 to 5 replicates. (G) Merging assays with aggregates of 75 cells size composed of E14 cells and TbxTKO cells. TbxTKO cells present arrested coalescence. Merging experiments taking place in a low adhesion U-bottom 96-well plate. N = 3 replicates.

During this time, gastruloids exhibit active cytoskeleton contraction from 96h (Figure 3B) as described for cells undergoing convergent-extension (reviewed in Sutherland et al, 2020). Furthermore, cells in gastruloids form actin enriched protrusions, and exhibit both filopodia- and lamellipodia-directed cell movement (Figure 4B) more actively in elongating regions (Figure 4A, movies 4 and 5). These protrusions have been associated with CE during axial elongation (reviewed in Wallingford et al, 2002; Wallingford et al, 2000; Belmonte et al, 2016) and are necessary for mouse mesoderm cells migration during gastrulation (Saykali et al, 2019). It may be that this activity is involved in axial elongation in gastruloids. In this context, we also observe the expression of the planar cell polarity (PCP) component *Wnt5a* in the posterior elongating region (Figure 4D). PCP is essential for CE (Wallingford et al, 2000; reviewed in Sutherland et al 2020) and *TbxT* mutant gastruloids do not elongate (Figure 4C, Wehmeyer et al., 2022) and their cells do not exhibit the protrusions.

**Figure 4.**
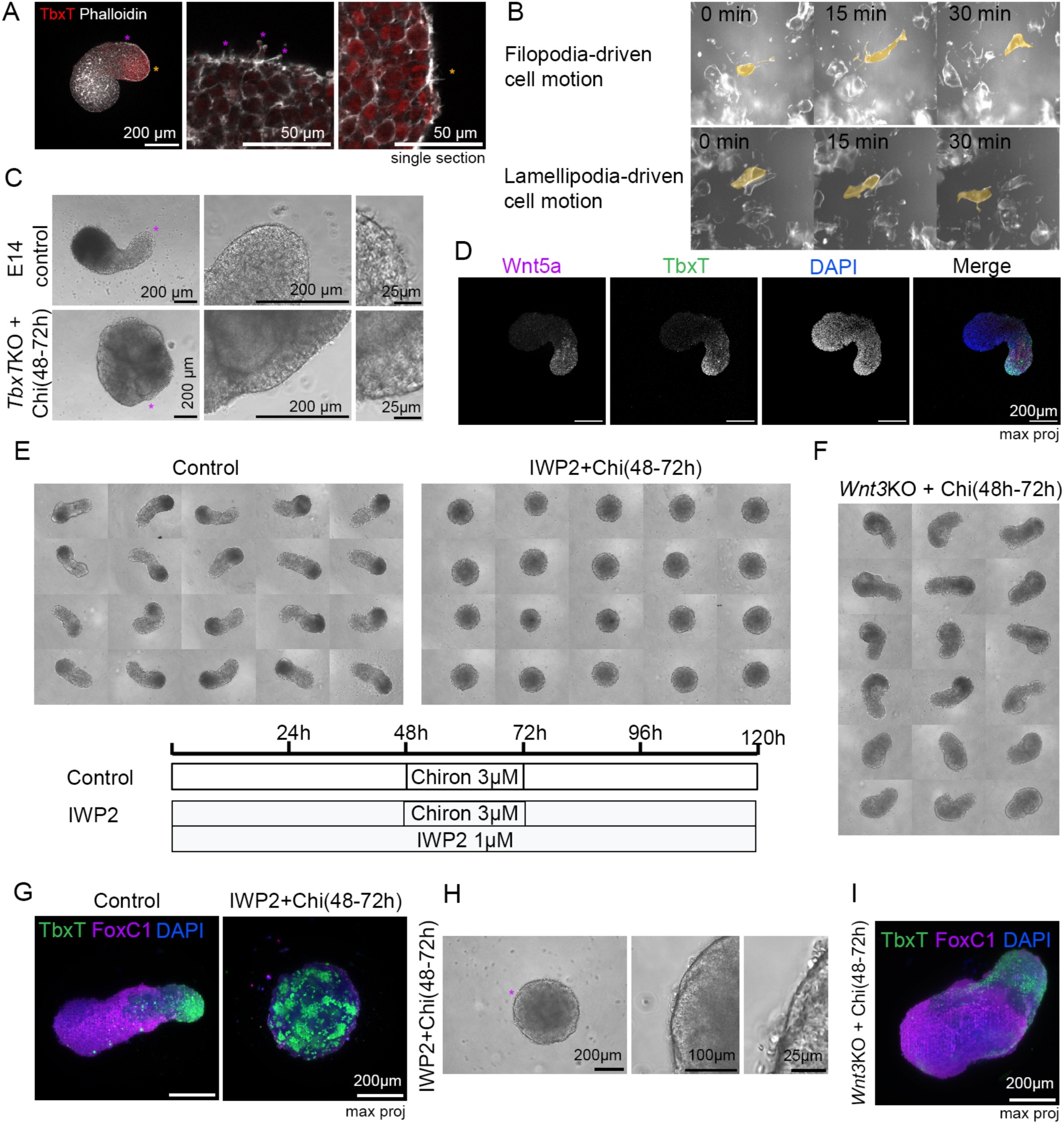
Wnt mediated PCP is necessary for gastruloid active protrusion formation and TbxT sorting during elongation. (A) Actin enriched filopodia formation in elongating gastruloids. (B) Types of cell movement (filopodia- and lamellipodia-driven cell motion) observed in chimeric gastruloids with membrane sparse-labelling (75% E14 cells + 25% R1(GPIGFP;H2BmCherry) cells) imaged with light sheet microscopy. (C) Brightfield images of E14 and TbxTKO 120h gastruloids highlighting differences in the amount of protrusions formed. (D) HCR labelling of TbxT and of the non-canonical Wnt molecule Wnt5a in 300-cell 120h gastruloids. (E) Brightfield images (and experimental scheme) of the effects of blocking non-canonical Wnt signalling on elongation. IWP2 prevents Wnt molecules secretion and Chiron (a Wnt agonist) rescues intracellularly canonical Wnt signalling. N = 3 replicates. (F) Brightfield images showing a pulse of 3 μM Chi between 48h-72h is enough to rescue elongation in Wnt3 mutant (Wnt3KO) gastruloids. N = 3 replicates. (G) Effects of IWP2 treatment on the expression pattern of expression of TbxT. N = 3 replicates. (H) Brightfield images of IWP2 (1μM) treated gastruloid without protrusions. N = 3 replicates. (I) Wnt3KO 120h gastruloid with 3μM Chiron pulse between 48h-72h exhibiting elongation, anterior expression of FoxC1 and posterior expression of TbxT. N = 3 replicates. Coloured text star symbols indicate magnified regions.

### A Tbxt:Wnt signalling axis in polarisation and elongation

*TbxT* has been shown to control the movement and directionality of cells during gastrulation (Hashimoto et al, 1987; Wilson and Beddington, 1997; Turner et al, 2014). It regulates the expression of members of both canonical and non-canonical Wnt signalling in mice (Evans et al, 2012; Su et al, 2024) and is itself regulated by Wnt signalling (Arnold et al, 1999, Yamagushi et al, 1999; Arnold et al, 2000). In the embryo, *TbxT* expression is initiated by Wnt3 (Tortelotte et al 2013) and, in agreement with this, *Wnt3* mutant gastruloids remain round (Figure S4B) and lack expression *TbxT* expression (Figure S4C). On the other hand, *TbxT* mutant gastruloids do not elongate and exhibit reduced *Wnt3a* expression at 96h and 120h (Wehmeyer et al, 2022), and localised *Wnt5a* expression probed by HCR at 120h (Figure S4A).

To further explore the relationship between *TbxT* and Wnt signalling in gastruloids, we separated canonical and non canonical signalling with combinations of agonists and antagonists of Wnt signalling. Treating gastruloids between 24-48h with IWP3, a pan-inhibitor of Wnt secretion, and providing CHIR between 48 and 72 hrs delays their elongation and *Tbxt* expression (Turner et al, 2017). However, gastruloids treated with IWP2 (another pan-inhibitor of Wnt secretion), from the time of aggregation, with addition of CHI between 48 to 72h, do not elongate (Figure 4E) suggesting that Wnt/ß-catenin signalling plays a role in the early stages of gastruloid development that is not rescued by CHIR. A closer look at the cells in these gastruloid shows that they lack the protrusions that we had observed under standard conditions (Figure 4H). These observations suggest that gastruloid elongation might be dependent on non-canonical Wnt signalling. This is supported by the observation that when *Wnt3* mutant gastruloids, which still are functional for non canonical Wnt signalling, are treated with CHIR, they recover both *TbxT* and elongation (Figures 4F and 4I). Furthermore, treating CHIR driven gastruloids with IWP2 from 48h or from 72h prevents elongation and adding Wnt3a from 72h does not rescue the IWP2 induced lack of elongation (Figure S5). Addition of CHIR between 48 and 72h to gastruloids treated with IWP2 from cell seeding, rescues the expression of *TbxT* which now is observed in a large number of foci dispersed throughout the gastruloids (Figure 4G). This is indicative of a role for non-canonical Wnt signalling in cell sorting.

The role of feedbacks between TbxT and Wnt signalling during polarization and elongation of gastruloids are reinforced by the observation that CHI does not induce elongation in *TbxT* mutant gastruloids (Figure 4C). This suggests that PCP is involved in this process downstream of Wnt signalling in coordination with TbxT. We then explored the role of other cellular properties on the dynamics of elongation and their relation to size.

### Size-dependent mechanisms of morphological dynamics

It is well established that tissue mechanical properties intrinsically influence morphogenesis (reviewed in Trubuil et al, 2021; Petridou and Heisenberg, 2019). As the patterns of cell division are fairly homogeneous in developing gastruloids, fluidisation of the tissue through cell divisions and differential growth, are unlikely to play a major role in gastruloid morphogenesis (Figure S6A). The correlation between the size of the initial gastruloid and the onset of elongation (see Figure 2A) suggests that gastruloid size might affect tissue mechanics. For this reason, we explored whether gastruloid size affects tissue mechanical properties using different strategies. To do this, we made use of the observation that between 72 and 96hrs, gastruloids have an ability to regulate their patterning upon dissociation. If gastruloids are dissociated into single cells and reassociated at 72 hours, they will polarize and elongate but this ability is lost by 96hrs (Figure S6B). This suggests that at 72hrs, cells in the gastruloid are not yet committed to a global pattern. Consistent with this, if we brought together two 72hrs aggregates of 20, 75 or 300-cell gastruloids each, one labelled with Hoechst and one unlabelled, they join in the middle and, with a size dependent dynamics, form a single round aggregate with cells mixed across boundaries before proceeding to a monoaxial gastruloid (Figure 3D-G; movies 7 and 8).

This experiment allows us to gain some insights into the mechanical properties of the gastruloids (Oriola et al, 2020) by quantifying the dynamics of aggregate merging across sizes. To do this we measured the initial radius of the aggregates and correlate it with the time it takes for the complete coalescence of gastruloids into a single spherical aggregate (assessed as the time at which the merging complex reaches maximum circularity in 2D; Figure 3D and 3F). Measurements of this process reveal a linear relationship between complete merge time and the initial radius of the aggregates (Figure 3F). Following a force balance mechanical model of aggregate fusion established by Oriola and colleagues (Figure 3D; Oriola et al, 2020), which predicts the speed of merging to be proportional to the initial radius of the aggregates (assuming equal mechanical properties across sizes), this is indicative that at 72h gastruloids have the same tissue mechanical properties independently of the size.

This behaviour parallels the relationship between *TbxT* expression, size and elongation that we have described above. To assess this further, we compared aggregation between *TbxT* mutant and between wild type gastruloids. We observe that the dynamics of aggregate merging between *TbxT* mutant cell aggregates with *TbxT* mutant cell aggregates is different from that of wild type aggregates with themselves, with *TbxT* mutant aggregates exhibiting arrested coalescence (29/29 with 75 cells aggregates and 27/27 with 150 cells aggregates; Figure 3G). WT and *TbxT* mutant gastruloids merged at 72h present mixed phenotypes (Figure S7). The majority of merged gastruloids develop a certain degree of mixing, with interspersed pure WT domains, and fail to elongate (13/20). A lower percentage of gastruloids keep the WT and *TbxT* mutant domains fully segregated and elongate (7/20). This supports the suggestion that an important function of *TbxT* is to change the mechanical properties of differentiating cells leading to an increase in tissue fluidity and movement (Wilson and Beddington, 1997; Turner et al, 2014).

Work from Gsell and colleagues (Gsell et al, 2024) points towards TbxT positive cells having a lower surface tension than other cells during gastruloid development. To test the role of cell surface tension during axial extension in gastruloids, we induced a reduction in actin polymerization, which should decrease cellular cortical tension, by applying a low dose of cytochalasin D between 90-120h (Fig. 3C). This leads to more prominent elongation (Figure 3C, movie 6).

Gastruloids form cellular protrusions during elongation (Figure 4A and 4B). *TbxT* mutant gastruloids exhibit a considerably lower amount of protrusions in comparison to wild type ones (Figure 4C). Treating gastruloids with a ROCK inhibitor (Y27632) has been reported to inhibit gastruloid elongation (Jong et al, 2023). This indicates that *TbxT* is necessary for establishing the correct cellular behaviour and mechanical properties that enable elongation. We then explored the relationship between multi axis formation and *TbxT* spatial expression dynamics.

### Gastruloid size effects on polarisation and elongation

While smaller gastruloids (< 70 cells) exhibit variable elongation, large gastruloids tend to form structures with two or more axes and the frequency increases with size (Figure 1D). As the pole of *TbxT* expression results from the coalescence of clusters of *TbxT* expression (Figure 5A), we wondered if this could be a result of Cadherin (Cdh) mediated cell sorting as suggested for endodermal tissue (Hashmi et al, 2022). For this reason, we first looked at the expression of E and N-Cdh during elongation.

**Figure 5.**
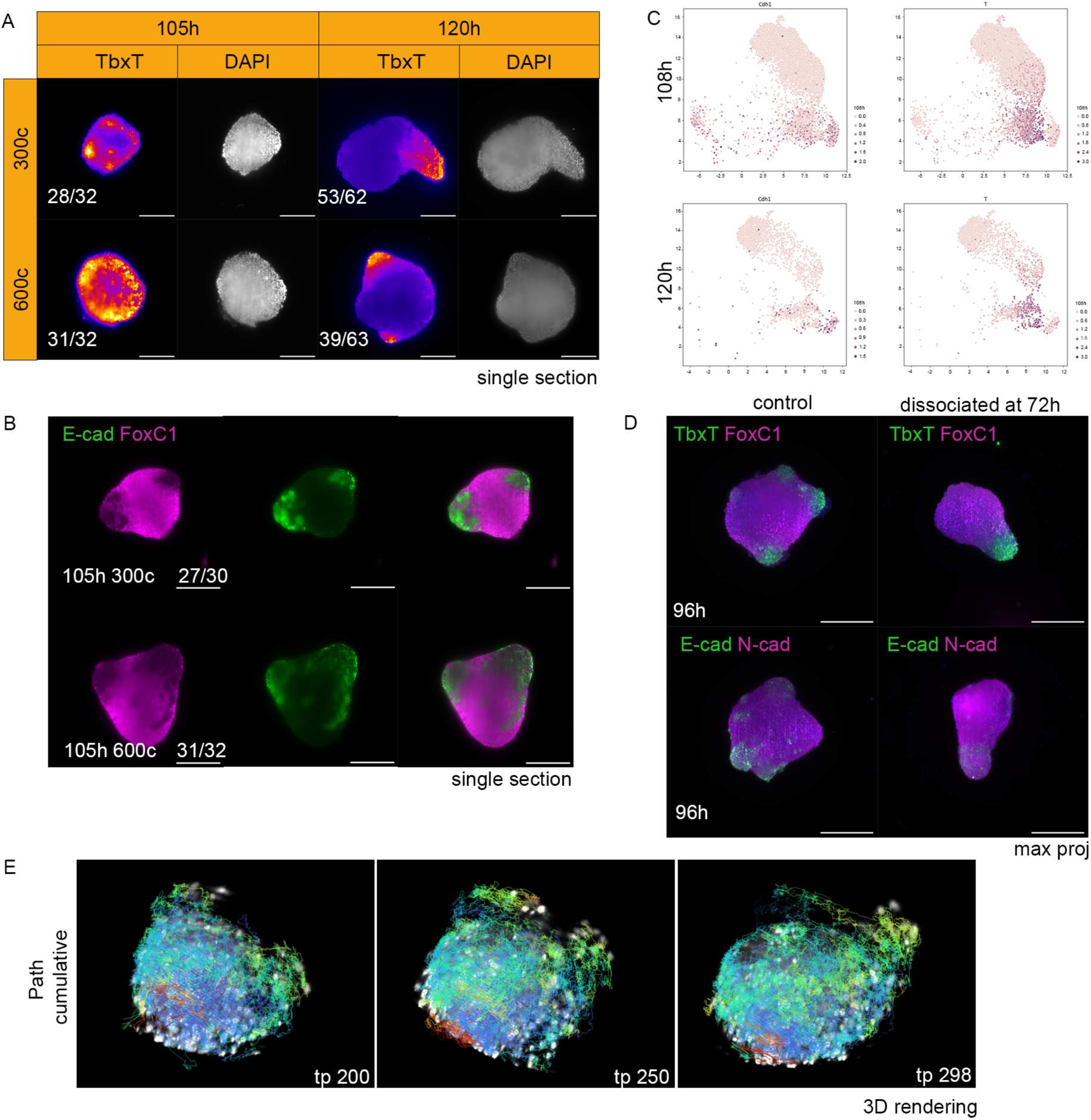
Large gastruloids multi-axis formation. (A) At 105h TbxT expression in 300 and 600-cell gastruloids is polarised into multiple foci. At 120h TbxT expression has been resolved to one single domain and single axis or alternatively has given rise to multiple axis formation and multiple domains. Light-sheet imaging of immunofluorescence, single section. N = 3 replicates. (B) At 105h E-cadherin expression is enriched in bulges. Light-sheet imaging of Immunofluorescence, single section. N = 3 replicates. (C) UMAPs for E-cadherin and TbxT expression at 108h, and 120h showing an overlap in expression. The UMAPs were produced using the publicly released data from the Liberali laboratory (Suppinger et al, 2023). Highlighted representation of E-Cadherin (Cdh1) and TbxT (T). (D) Dissociating and reassociating gastruloids at 72h speeds up gastruloid development: at 96h both TbxT and E-cadherin expression are polarised and gastruloids have elongated. Light-sheet imaging of immunofluorescences, max projection. (E) Nuclear movement with converging tissue bulges during elongation. Light sheet microscopy data derived from a chimeric gastruloid with nuclear sparse-labelling (60% E14 cells + 40% emiRFP670 cells). Scale bar – 200μm.

Previous to the Cadherin switch (Figure S8), the *TbxT* expressing cells also express E-Cdh (Figures S9 and S10). This situation is reminiscent of that in the embryo where in the early primitive streak E-Cdh expression overlaps with *TbxT* (see Fig S11). In agreement with previous reports, we also observe a temporal correlation between elongation and a switch from E-to N-Cdh (Figure S9 and S10) that can also be observed during gastrulation (Figure 5, S11 and S12). These regional Cdh expression patterns are likely to regulate the dynamics of the coalescence of protruding morphological deformations of *TbxT* multi foci, and help establish the elongating axis/axes (Figure 5B, 5D, S8-S11, movie9).

The results are consistent with the timing of elongation initiation and *TbxT* multifoci coalescence being independently regulated. In a big proportion of large gastruloids multi-foci do not resolve into a single domain prior to the onset of elongation, resulting in the formation of multi-axes. This highlights the importance of size regulation during embryonic development for ensuring correct embryo morphologies. We observe that in experiments where gastruloids were dissociated and reassociated at 72h, the morphogenetic process of gastruloid elongation and cell sorting occurred considerably faster (Figure 5D). This can be construed as lower adhesion properties accelerating morphogenesis and cell sorting. In gastruloids forming multi-axis we observe that cells in bulging regions move towards neighbouring bulges (Figure 5E and movie9).

When analysing the relation between the number of axes and gastruloid size within multi-axial gastruloid samples, we observe that larger gastruloids present globally a higher number of axes (Figure 1, Figure S13A). Normalised angles between contiguous axes (only angles smaller than 360 divided by the number of axes were analysed) have a wide distribution but are significantly bigger in 120h, 600-cell gastruloids than in 300-cell gastruloids (Figure S13B).

### Size-dependent divergences in gene expression and cell fate specification outside robust size range

While large gastruloid sizes can lead to multi-axes formation, very small gastruloids at times fail to elongate (Figure 1A). This different behaviour led us to search for differences in gene expression in these structures. Although globally all gastruloids seem to have similar gene expression patterns (Figure 1E-F), comparison between the two extreme sizes of our experiments (20 and 300 cells), highlight differences in terms of DGE (Figure S14). The differences are mild but significant.

When compared to 300-cell gastruloids, 20-cell gastruloids exhibit a down-regulation of genes related to hypoxia and metabolism (Figure S15A) and an upregulation of genes related to neural development (Figure S15B). This can be seen in spatial gene expression as 20-cell gastruloids have significantly higher expression of neural marker genes such as *Sox1*, *Sox2* and *Sox3* (Figure S15C) and a downregulation of paraxial mesoderm genes such as *Tbx6* and *Msgn1* (Figure 6A). *Tbx6* is not detected by immunofluorescence in most (but not all) 20-cell gastruloids and always present in 300-cell gastruloids, whilst Sox2 and Sox3 expression is clearly present in both 20 and 300-cell gastruloids at 120h (Figure 6D-E and S15C). There is thus a differential relative tissue composition.

**Figure 6.**
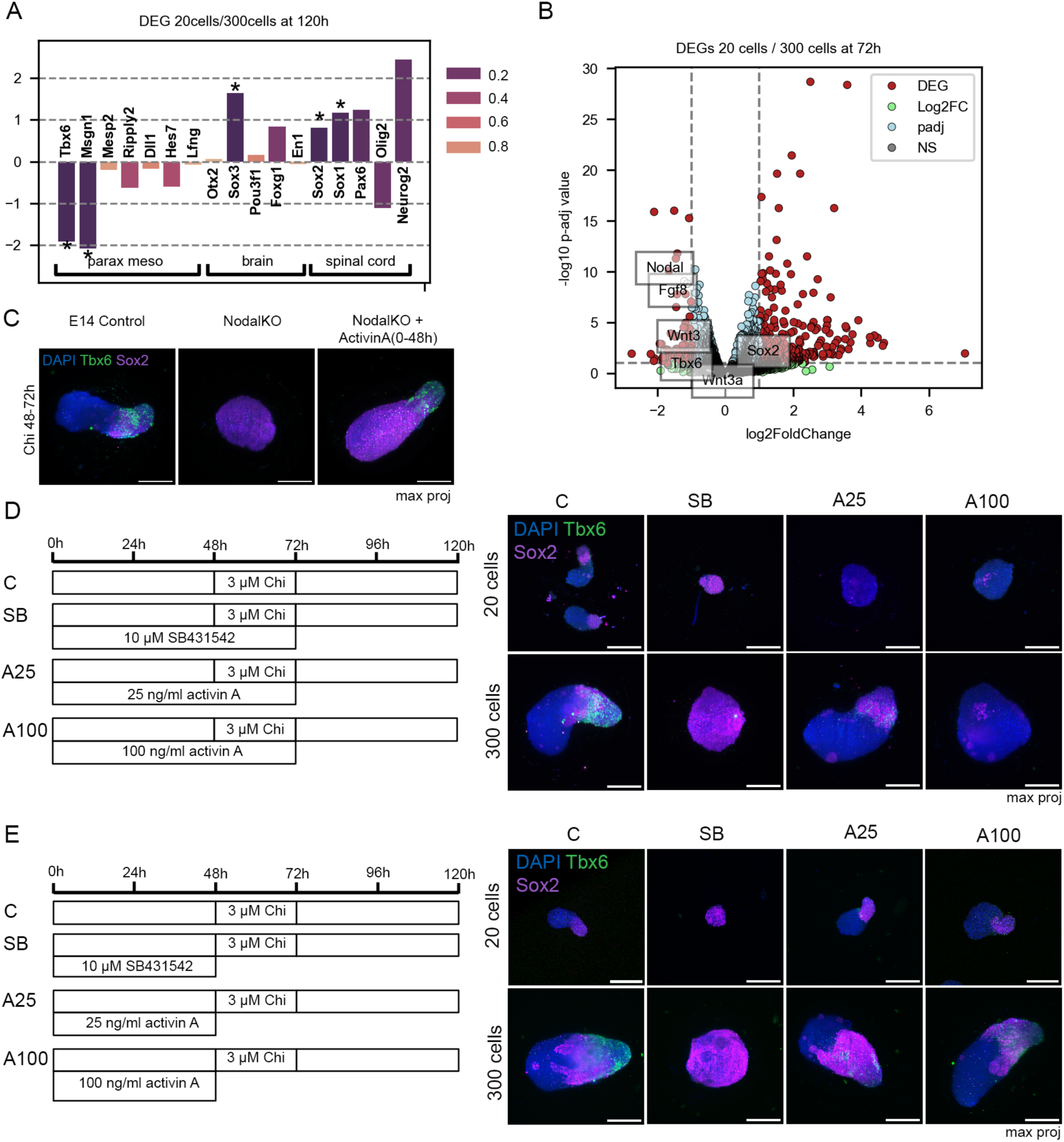
Smaller gastruloids have lower Nodal expression which modulates neural and paraxial mesoderm fates specification. (A) Differential gene expression of fate markers between 120h 20-cell and 300-cell gastruloids. (B) Differential gene expression volcano plot comparing 20-cell and 300-cell, 120h gastruloids. DEGs thresholds were set at LFC>1 and p-adjusted value<0.1. mRNAseq analysis from bulk mRNA with N = 3 replicates. (C) Analysis of the patterns of Sox2 and Tbx6 expression in 300-cell *Nodal*KO 120h gastruloids and in 300-cell *Nodal*KO 120h gastruloids rescued by a 0-48h treatment with 25ng/ml ActivinA treatment. N = 3 replicates. (D) Experimental scheme and effects on Sox2 (neural marker) and Tbx6 (paraxial mesoderm marker) expression, of 3 days perturbation of Nodal signalling activity in 120h gastruloids. N = 3 replicates. (E) Experimental scheme and effects on Sox2 and Tbx6 expression, of 2 days perturbation of Nodal signalling activity in 120h gastruloids. In the conditions A25 and A100 for 20-cell gastruloids not all samples present Tbx6 expression. N = 3 replicates. Scale bar 200μm.

We also notice that, relative to 300-cell gastruloids, 20-cell gastruloids present a downregulation of the expression of *Nodal,* as well as of its targets *Lefty1* and *Lefty2* expression at 72h (Figure 6B and S16). Nodal signalling is known to inhibit neural development (Camus et al, 2006) and this led us to test whether differences in Nodal signalling activity between the two sizes could explain the differential tissue composition (Figure 6C-E).

Loss of function of Nodal signalling activity through exposure to the Alk4/5/7 receptor inhibitor SB431542, results in a very marked increase in *Sox2* expression and prevents *Tbx6* expression in 20-cell and 300-cell gastruloids (Figure 6D). On the other hand, an increase in Nodal signalling activity with a 0-72h Activin treatment reduced Sox2 expression in a concentration dependent manner (Figure 6D). We notice that *Tbx6* expression is downregulated with a 0-72h Activin treatment, while a 0-48h Activin treatment does not have noticeable effects in *Tbx6* expression at 120h in 300-cell gastruloids and Tbx6 expression can be seen in some 120h 20-cell gastruloids (Figure 6E). In agreement with these assays, *Nodal* mutant gastruloids have very strong *Sox2* expression, do not express *Tbx6* and do not elongate. Providing Activin to *Nodal* KO gastruloids during the first 48h (with CHIR treatment at 48-72h) rescues elongation and *Tbx6* expression (Figure 6C). Nodal signalling downregulates *Sox2* expression and Nodal activity is required during the first 48h of gastruloid development for *Tbx6* expression (Figure 6C-E).

## Discussion

Size control and pattern scaling are intrinsic elements of animal embryogenesis. Here, using PSC derived gastruloids we were able to explore the effects of size perturbations on morphogenesis. Gastruloids allow the experimental exploration of several features of mammalian gastrulation in the absence of many of the constrains associated with the preimplantation development, in particular, the interactions between extraembryonic and embryonic cells.

A conspicuous feature of gastruloids is the elongation of the initial aggregate that starts between 72 and 96 hours after aggregation upon exposure to the Wnt agonist Chiron. This event can be associated with axial elongation in the embryo. However, comparison of the patterns of gene expression between gastruloids and embryos (Suppinger et al, 2023; Sala et al, 2019 and Figures S8-12) lead us to suggest that this event corresponds to the late gastrulation phase in embryos, around E7.5 and E8.0. We surmise that the initial elongation observed in gastruloids corresponds to the reorganisation that in the mouse embryo takes place between E7.5 and E8.0 that gives rise to a recognizable vertebrate body plan. During this time the characteristic cup shape of the early mouse embryo is transformed into an elongated structure with the three basic axes (Fig. S17) and, in the posterior part, the formation of the Caudal Lateral Epiblast. There is some evidence that this transformation is associated with Planar Cell Polarity (PCP) signalling (Mahaffey et al., 2013). This correlation allows us to interpret the main findings of our study and explore what they might tell us about events in the embryo.

Here we have exploited the ability of gastruloids to form and scale within a range of starting cell numbers to study the effect of size on morphogenesis. Our results allow us to extract a number of conclusions about the development of the gastruloids that can be extrapolated to the embryo.

### (1) TbxT fluidifies the mesodermal tissue enabling elongation

We observe a correlation between the onset of *TbxT* polarisation and gastruloid elongation. In the absence of *TbxT* there is no elongation and smaller gastruloids that exhibit an earlier polarisation of *TbxT* expression, elongate earlier (Figure 2). Furthermore, our experiments merging aggregates suggest that *TbxT* expression enables a fluid-like tissue behaviour, a factor that may be essential for tissue elongation (Petridou and Heisenberg, 2019). In gastruloids with a mixed composition of WT and *TbxT* mutant cells, while WT cells disperse randomly, *TbxT* mutant cells exhibit a sorting behaviour (Wehmeyer et al., 2022). This further supports different tissue mechanical properties between the two genetic backgrounds. Merging WT and *TbxT* mutant gastruloids presents partial WT/TbxTKO cell domains segregation (Figure S7), suggestive of different tissue mechanical or adhesive properties. Work from Gsell and colleagues (Gsell et al, 2024) suggests that TbxT positive cells have lower surface tension than TbxT negative cells which might be responsible for this behaviour. In agreement with this suggestion, mildly treating gastruloids with cytochalasin D (an inhibitor of Actin polymerisation) results in increased elongation (Figure 3C).

The TbxT mediated tissue fluidity is likely to be related to the observation that TbxT mediates autonomous cell movement into and within the primitive streak as well as in tissue culture and there is evidence that this movement could be directional (Turner et al, 2014; Kwan and Kirschner, 2003, Wilson and Beddington, 1997). It is possible that, at the ensemble level, this movement could result in the global tissue elongation that we observe. Our light sheet cell movement analysis indicates that the posterior elongating region of gastruloids has strongly higher cell motility and mildly higher directional cell movement when compared to the anterior region (Figure S2B). However, the source of the global directionality, particularly in the gastruloids, is not clear. In the embryo it is likely that this is imposed by a coordinate system created by the asymmetric organization of the extraembryonic tissues.

### (2) Wnt-mediated PCP is needed for gastruloid elongation, not only because of its role in directing cell movement but also for the sorting of TbxT expressing cells

During gastruloid development, we observe that foci of *TbxT* expressing cells arise independently and coalesce to establish discrete domains defining the AP axis. Elongation initiates without requiring a complete coalescence of TbxT expression domains indicating the onset of the elongation programme is under some global influence independent of full TbxT polarisation. We have begun to investigate the mechanisms driving these events. In principle, at the global level the extension in gastruloids is compatible with a Convergent Extension (CE) movement (de Jong et al, 2024).

The elongation is associated with coalescence of TbxT expressing cells in a manner reminiscent of what has been observed before in gastruloids treated with Activin and CHIR (Hashmi et al, 2022). However, as Activin treated gastruloids do not elongate (van den Brink et al, 2014; Turner et al, 2017), it must be Wnt signalling that drives the elongation. This suggests an involvement of PCP through non canonical Wnt signalling.

There is evidence of a correlation between cell autonomous protrusion activity, PCP signalling within cell ensembles and CE in vertebrate embryos (reviewed in Davey and Moens, 2017; Wallingford et al, 2002). We observe many of these elements associated with gastruloid elongation. For example, we observe protrusion activity in single cells within the elongation domain of the gastruloid that is reduced when we block Wnt/PCP signalling (Figure 4H). In these gastruloids, there are still *TbxT* expressing cells that fail to coalesce into a single pole suggesting that Wnt/PCP signalling is not only involved in global tissue behaviour, but also in driving directional coalescence of individual cells. It is likely that TbxT contributes to this process by controlling not only Wnt signalling but also components of cytoskeletal regulatory networks (Su et al, 2024).

Thus, it appears that Wnt-mediated PCP plays an important role in mediating cell sorting and tissue patterning in developing gastruloids. However, whether CE is involved in this process is not clear from our results. An earlier study has suggested that gastruloids undergo CE through active cell crawling and differential adhesion on the basis of morphological analysis, Cellular Potts modelling simulations and limited manual cell tracking (de Jong et al, 2024). The reported tracking results present directional movement but do not exhibit unambiguous features of convergent extension movements. Our results show high cell motility with a net posterior movement, but do not reveal clear evidence for CE either. This behaviour is consistent with studies of gastrulation in mouse embryos that have also failed to detect evidence of Convergent Extension guiding this process (Williams et al, 2011).

The importance of the dynamics of cell sorting in gastruloid self-emergent organisation has recently been highlighted also in the context of establishing signalling centres in the work of McNamara and colleagues (MacNamara, 2024). Given the cross transcriptional regulation between Wnt and TbxT, this hints at a Wnt-mediate PCP being a central organising factor in gastruloid development. The source of the polarised global behaviour of the tissue remains an open question.

### (3) The pattern of Cadherin molecules affects the dynamics of cell sorting and axis formation

Monopolar gastruloids only form within a certain range of cell numbers that suggests the existence of some global system that determines a range of cell diameters over which coalescence might occur. Small initial cell numbers lead to gastruloids in a stochastic manner, whereas an initial cell number of about 400 cells, leads to gastruloids with multiple axes (Figures 1 and 5). In general, we observe that the single pole of *TbxT* expression associated with elongation results from the coalescence of several foci of *TbxT* expressing cells. Some of these are enriched in E-cadherin. It may be that the coalescence is driven by differential adhesion. In support of this possibility, when dissociating and reassociating gastruloids (Figure 5D), hence drastically reducing adhesion, E-Cadherin+ cells are found to coalesce into a single pole, faster than in the control situation. This is reminiscent of the fact that gastruloids that polarise TbxT earlier (which fluidizes tissue, like lower adhesion) initiate elongation earlier. Another example of adhesion mediating gastruloid elongation is the observation that *N-Cdh* mutant gastruloids exhibit increased elongation (Mayran et al, 2023).

The notion that E-Cdh contributes to coalescence of cells in gastruloids is supported by reports on the formation of endoderm in gastruloids (Hashmi et al, 2022; Vianello and Lutolf, 2021). This also might correlate with a transient state during mid and late gastrulation in the embryo, when TbxT expressing cells also express E-Cdh (Figure S11).

These observations also suggest an explanation for the emergence of multiple axes. If the initial aggregate is too large, elongation might ensue before the coalescence into a single pole (Figure 7).

**Figure 7.**
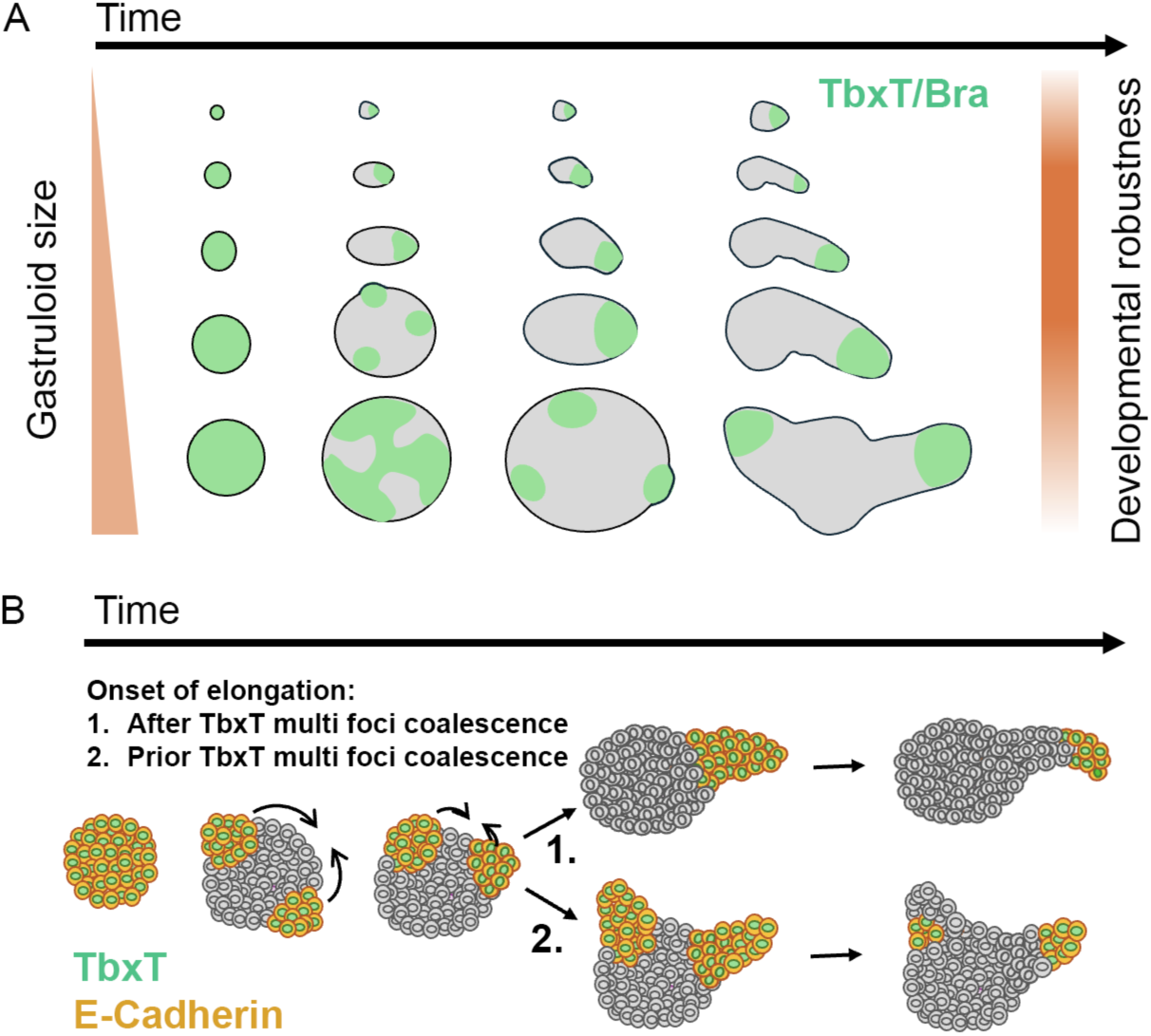
Size regulates gastruloid axis formation and axial elongation through TbxT polarisation. (A) Dynamics of TbxT patterning across sizes during gastruloid development. There is a robust size range for the formation of monoaxially elongated gastruloids. Smaller gastruloids initiate elongation and TbxT polarisation earlier. TbxT expression evolves from homogeneous expression, to patched expression, to discrete domains that coalesce into the future elongating region(s). Large gastruloids can form multi-axes. (B) Temporal independence of TbxT positive cells sorting and onset of axial elongation during gastruloid axis formation. TbxT/E-Cdh multi-foci coalesce into the domains where elongation will take place. Differential cell adhesion likely modulates these dynamics. If elongation begins prior to full coalescence the gastruloid will establish multi-axes.

### (4) The size of the initial aggregate impacts the dynamics of the elongation and the cell fates in the gastruloid

Smaller gastruloids can present morphological defects and lack of proper elongation, and from bulk RNAseq analysis they seem to favour neural development, having low expression of genes from paraxial mesoderm such as *Tbx6*. We explain this through a difference in Nodal gene expression. During mouse embryonic development, Nodal signalling is required for the formation and maintenance of the primitive streak (Colon et al, 1994), and furthermore blocks early neural differentiation (Camus et al, 2006). In small gastruloids, which present downregulated Nodal signalling activity, we see this behaviour being recapitulated with increased expression of neural markers and concomitant loss of some mesodermal tissue (paraxial mesoderm). Nodal activity is necessary during the first 48h of gastruloid development for the formation of paraxial mesoderm. Providing Activin A (25ng/ml) to Nodal mutant gastruloids during the first 48h of gastruloid development is enough to rescue Tbx6 expression (paraxial mesoderm marker) and the gastruloid elongated morphology.

In summary, here we report that in gastruloids TbxT elicits fluidisation and directional movement of cell ensembles, some of which is driven by differential adhesion and some by Wnt/PCP. These cell movements are important for defining tissue spatial organisation, signalling centres and for morphogenetic anteroposterior axial elongation. These observations build on the functional implications of the role of TbxT in regulating embryonic cell movement during mouse gastrulation (Wilson and Beddington, 1997) without the constrains of the embryo. In the embryo, the axial orientation imposed in the epiblast by the extraembryonic tissues is likely to determine cell movements in the primitive streak during gastrulation. In gastruloids, cells have a global directional movement whose source remains to be explored. This movement might be the substrate for the situation that is observed in vivo.

Our results also reveal a relationship between size and cell movement that might have implications in animal body axis formation. Size is an important variable in the morphogenetic movements as it might define the boundary conditions that set up the distances and timing of coalescing TbxT expression domains that will establish AP axes. Our observations also suggest that a minimal epiblast size is required for robust patterns of signalling activity and consequently of tissue fate specification. TbxT modulates tissue mechanical properties paramount in cellular sorting, morphogenetic movements and the establishment of patterning cues (signalling centres).

Overall, our results support the notion that large scale tissue movements play an important role in the organization of signalling centres and their impact on morphogenesis (Busby and Steventon, 2021; Fulton et al, 2020).

## Material and Methods

### Plasmids

For producing E14-TG2a Nodal KO and E14-TG2a Wnt3 KO lines, pSpCas9(BB)-2a-GFP-gRNA plasmids were generated.

Nodal KO guide RNA (gRNA) oligo sequences containing flanking *BbsI* restriction sites:

gRNA1_F: 5’-CACCGCGTGAAAGTCCAGTTCTGTC-3’

gRNA1_R:5’-AAACAGACAGAACTGGACTTTCACGC-3’

gRNA2_F:5’-CACCGGTACAATGCCTATCGCTGTG-3’

gRNA2_R:5’-AAACACACAGCGATAGGCATTGTACC-3’

Wnt3 KO guide RNA oligo sequences containing flanking *BbsI* restriction sites:

gRNA1_F: 5’-CACCGGTACCTGCTGGCCCAGGGCCA-3’

gRNA1_R:5’-AAACATGGCCCTGGGCCAGCAGTACC-3’

gRNA2_F:5’-CACCGGGCCATCTTTGGGCCTGTCT-3’

gRNA2_R:5’-AAACAGACAGGCCCAAAGATGGCCC-3’

The gRNA oligos were annealed with a cycle of 94C for 4 min, 70C for 10 min and 37C for 20 min and introduced into the pSpCas9(BB)-2A-GFP plasmid (Addgene, #48138; Ran et al, 2013) *BbsI* sites with Golden Gate assembly.

### Cell culture

The cell lines used were E14-TG2a (50-238-2109; ATCC), H2B-mCherry;GPI-GFP (R1 ES cells; Novotchin et al, 2009), Bra-/-mT (Schüle et al, 2023) and Bra-GFP;Sox17-RRFP (Pour et al, 2022). E14-TG2a ES cells were used for generating a Nodal knock out (KO) and a Wnt3 KO transgenic line. The new mouse transgenic ES lines were generated by CRISPR-Cas9 deletion using 2 single guide RNAs targeting respectively, exon 2 of the *Nodal* gene (ENSG00000156574; Nodal KO line), and targeting the 5’ and the 3’ region of exon 2 of the *Wnt3* gene (ENSMUSG00000000125; Wnt3 KO line). Expression of Cas9 and guide RNAs (gRNA) was done by lipofectamine transfection (Lipofectamin 3000; Thermo Fisher Scientific; L3000001) of a pSpCas9(BB)-2a-GFP-gRNA plasmid with single cell clonal line GFP positive sorting 1 week after transfection and knockout validation by PCR-based genotyping of amplified clones for the Nodal KO line and by reverse transcription-quantitative polymerase chain reaction (RT-qPCR) for the Wnt3 KO line.

Nodal test amplification primers:

5’-GCAGGTTCCAATTCTGCCTC-3’

5’-GAGCGCCCTCTCCCAATG-3’

Wnt test amplification primers:

5’-CCGTGCCCCAAGGAAAGTC-3’

5’-ACTGTGTCATCCGGGAAGGA-3’

Lastly the selected clones were validated by sequencing (Figures S18 and S19).

E14-TG2a and H2B-mCherry;GPI-GFP pluripotent stem cells were grown in ESLIF media (500 ml G-MEM BHK-21 (Gibco; 21710025), 5 ml Sodium Pyruvate (100mM; Gibco; 11360-039), 5 ml non-essential amino acids (Gibco; 11140-035), 5 ml GlutaMAX (Gibco; 35050-061), 5 ml PennStrep (Gibco; 15070-063), 1 ml b-mercaptoethanol (50 mM in PBS), 550ul of 10ug/ml LIF (Qkine; Qk018)) with 10% FBS (Gibco; Ref 10270-106) and Bra-GFP;Sox17-RRFP in ESLIF media with 15% FBS (Gibco; Ref 10270-106), changing media every day. Confluent pluripotent stem cell cultures were passaged with Trypsin-EDTA (0.5%), phenol-red (25300-054; ThermoFisher Scientific).

### Chemical reagents

The chemical reagents used were cytochalasin D (C2618-200UL; Sigma-Aldrich; 0.05 ng/ml, 0.5 ng/ml), Activin A (Qk005-500ug; Qkine; 25ng/ml, 100ng/ml), Chiron (SML1046-5MG; Sigma-Aldrich; 3mM), SB431542 (1614; Tocris; 10μM), Hoechst 33342 (Fisher Scientific; 62249; 1:40000), DAPI (ThermoFisher Scientific; D1306; 1:3000), Phalloidin-iFluor647 (Phalloidin-iFluor 647; ab176759; Abcam; 1:4000), IWP2 (Merck Life Science/SIGMA; I0536-5MG; 1μM), Wnt3a (315-20; Life Technologies S.A.; 100ng/ml). Pharmacological treatments were performed as described in the figures.

### Gastruloid work

mESC derived gastruloids were prepared as previously described (van den Brink et al, 2014; Baillie-Johnson et al, 2015). In the dissociation/reassociation experiments, gastruloids were pooled at the desired time, rinsed once with warm PBS (calcium free). Gastruloids were dissociated by incubating in 1ml accutase at 37C for 5 min. Gastruloids were then fully dissociated by pipetting up and down 15 times and single cells recovered by centrifugation (200g for 3min). The cell pellet was resuspended in 240 microliters of N2B27. Cells from dissociated gastruloids were seeded in low-adhesion U-bottom 96 wells plate at 3000 cells per 150 microliter per well for gastruloids dissociated at 72h AA and at 6000 cells per 150 microliter per well for gastruloids dissociated at 96h AA. The plated cells were centrifuged at 150g for 2 min and then stored in the incubator.

### Immunohistochemistry and HCR

The primary antibodies used were goat anti-Bra (AF2085; R&D Systems; 1:100), goat anti-E-Cadherin (AF648; R&D Systems; 1:500), rabbit anti-N-Cadherin (ab18203; Abcam; 1:200), rabbit anti-Sox2 (ab92494, Abcam; 1:200), rabbit anti-Sox3 (PA5-35983; Thermofisher), goat anti-Tbx6 (AF4744; R&D Systems; 1:200), rabbit anti-PhosphoMyosin Light Chain 2 (Ser19; 1PMyosin) (3671S; Cell Signalling; 1:50), rabbit anti-PhosphoMyosin Light Chain 2 (Thr18/Ser19; 2PMyosin) (3674S, Cell Signalling; 1:75). The secondary antibodies used were donkey anti-rabbit Alexa Fluor 647 (A31573; ThermoFisher; 1:500), donkey anti-Rabbit Alexa555 (ab150062; abcam; 1:500), donkey anti-goat Alexa Fluor 488 (A11055; ThermoFisher; 1:500), donkey anti-goat Alexa Fluor 647 (A21447; invitrogen; 1:500).

Immunofluorescences (IF) were done on samples fixed in 4% paraformaldehyde (PFA) in Phosphate Buffer Solution (PBS; D87537-500ML; Sigma-Aldrich), at 4C, overnight. In IF labelling actin enriched protrusions, gastruloid samples were fixed with 8%PFA with 1 unit per ml Phalloidin (to stabilise actin structures) all diluted in PBS, at room temperature for 30 minutes. Fixed samples were washed once in PBST (0.1% Triton in PBS), 2 times in PBS, blocked for 1h30min in blocking solution (2%BSA in PBST) and incubated overnight at 4C with primary antibodies diluted in blocking solution. Then, the samples were washed 6 times for 10 minutes in blocking solution and incubated overnight at 4C with secondary antibodies diluted in blocking solution and DAPI (D1306; ThermoFisher; 1:3000). Lastly, the stained samples were washed 6 times for 10 minutes in PBST and mounted for imaging. For phosphomyosin IF, fixed samples were washed 3 times for 5 minutes in PBS, blocked in blocking solution (10%FBS, 0.2%BSA, 0.2% Triton in PBS) for 1 hour, washed 3 times 5 minutes in PBS and incubated ON with primary antibodies (anti-1PMyosin or anti-2PMyosin) in solution B. The samples were washed 3 times for 10 minutes in PBS and incubated with secondary antibodies in solution A, at room temperature for 2h. Then washed 3 times for 10 minutes with PBS and incubated overnight with DAPI at 4C. Lastly, they were washed 3 times for 5 minutes with PBS and mounted.

Gastruloid HCR mRNA labelling was done as described in Henessy et al, 2023. HCR probes and amplifier hairpins were purchased from Molecular Instruments.

### Microscopy

Wide-field time-lapse movies of developing gastruloids were acquired using a Zeiss Axio Observer Z1 fluorescence microscope in a humidified CO_2_ incubator (5% CO_2_, 37°C). Confocal imaging was done in a Zeiss LSM980 microscope and in inverted and upright Leica SP5 microscopes. Light sheet microscopy imaging was performed in a Viventis LS1 microscope in a humidified CO_2_ incubator (5% CO_2_, 37°C). Light sheet timelapse images were acquired every 5 min and z-sections had a spacing of 2 microns between sections. Sample datasets of the light sheet time lapse imaging will be released in FigShare upon publication.

### Image analysis

Morphological analysis of brightfield images was performed using Ilastik (Berg et al., 2019) to generate segmentation masks and the python library scikit-image to extract morphological properties.

Cell motion analysis is performed using a custom pipeline. The process can be subdivided in preprocessing, track computation and track analysis. A diagram of the pipeline can be seen in Figure S20.

During preprocessing, we remove cellular debris from the plate by cropping a box around the gastruloid image to remove confounder artifacts for the next steps of the analysis. In addition, during imaging time, debris is accumulated over the well and around the gastruloid. We reduce the impact of this artifact by defining a parabolic mask that mimics the shape of the culturing well. Pixels under the parabola are set to zero. The center and curvature of the parabola are manually set for each gastruloid. Both of these steps have been performed with the help of a custom napari-widget.

During processing, we perform rigid registration to correct the global motion of the gastruloids. After this step, the output of the registration is cropped with a second bounding box for storage efficiency. To extract information of the deformations during gastruloid development, we perform non-linear registration over the rigid-registered images. Rigid and non-linear registration have been computed using the block-matching algorithm as implemented in the python wrapper vt-python (https://gitlab.inria.fr/morpheme/vt-python). Since the non-linear transformation objects from these images are very heavy, we have developed a compressed version of the same based on the K-NN regressor. A grid of points is created at pixel and each time over the region of the registered images above a threshold of 200 (with respect to a uint16 format of the images). Then the non-linear transformation is applied independently to the grid of points at each time in backwards direction (ecc. the transformation moves the points according to the non-linear transformation between times t to t-1). The set of original points plus the non-linear backwards-transformed new positions are used as regressors and predictors, respectively, for the K-NN regressor. The fitted predictors are then stored and used for the subsequent analysis. The K-NN predictor objects are of much smaller size than the non-linear transformations because not all the points of the image are stored due to the mask, achieving a massive size reduction. The K-NN has been implemented using the python version scikit-learn (Pedrosa et al., 2011). Finally, we use seeding points at the later time point of each recorded gastruloid and apply the K-NN backwards predictors recursively to obtain non-linear registered tracks. The seeding points used are the centroids of automatically segmented cells at the last time point. Cellular segmentation is performed using Cellpose (Stringer et al. 2021) in 2D using the “cyto3” model and z-stitching with an overlap threshold of 0.4 to generate 3D masks.

Finally, metrics for visualisation of cell motion and tissue deformation are computed using the tracks obtained. To quantify the amount of movement we use two metrics: cumulative path and diffusion of the cells. To quantify directionality of the movement we use drift. The cumulative path is computed as:

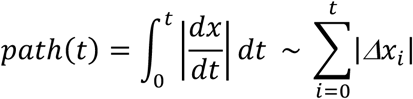

drift and diffusion are defined from the linear drift-diffusion equation:

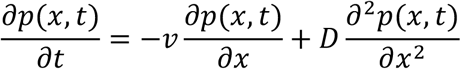

where *v* is the drift parameter and *D* is the diffusion parameter. Similar approaches have been used to characterize diffusion properties in colloid systems (Wieser et al, 2008). Which can be solved for a point initial conditions,

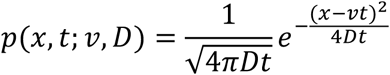

Using Bayes’ rule and considering uniform (improper) non-informative prior distributions, we can estimate the drift and diffusion parameters with its errors.

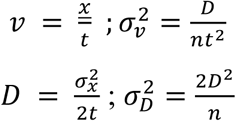

Where *n* is the number of samples we have per trajectory. We set the time to be 1, as the adimensional time between recorded images (5 min in natural time). We further make the approximation that the drift and diffusion are approximately constant during a small time window of the trajectories. We compute the metrics for each trajectory in windows of 5 steps (25 minutes, *n* = 5). The different metrics are used to color code the trajectories using a custom napari-widget.

### Bulk mRNA-seq of gastruloids

Gastruloid samples of different sizes (20 cells, 75 cells, 150 cells and 300 cells) were collected for total mRNA extraction at 48h, 72h and 120h after cell seeding. Gastruloids were collected into low binding eppendorf tubes washed twice with PBS (-) and mRNA was extracted using the Qiagen RNeasy Mini Kit following the manufacturer’s instructions. RNA-seq libraries of 120h gastruloids samples were prepared at CNAG (National Centre of Genomic Analysis, Barcelona, Spain) with polyA-enrichment, 100 bps paired-end reads and Agilent 2100 Bioanalyzer quality control, and sequencing with a 50M depth in an Illumina NovaSeq 6000 S1 system. RNA-seq libraries of the remaining gastruloids samples were prepared at the Genomics Unit of the IGC (Gulbenkian Institute for Science, Oeiras, Portugal) with polyA-enrichment, 100bp paired end reads and Fragment Analyzer (Agilent) quality control, and sequenced with an Illumina NextSeq 2000 system with a 25M reads depth. The sequencing data will be publicly released in the BioStudies database (formerly EMBL-EBI ArrayExpress).

### Bulk mRNAseq analysis

#### Data Preprocessing

The quality of the raw sequencing files was initially assessed using FastQC (version 0.11.9) software(Andrews, 2010). Adapter trimming and low-quality filtering (removing reads with a Phred score below 20) were performed using TrimGalore! (version 0.6.7)(Felix Krueger, https://github.com/FelixKrueger/TrimGalor). Following trimming, the quality of the cleaned reads was re-evaluated with FastQC as a quality checkpoint to ensure the effectiveness of the trimming and filtering steps.

The processed sequencing files were mapped to the mouse genome (GRCm38.p6) using the Gencode annotation (version 25) with the STAR alignment algorithm (version 2.7.10a) (Dobin et al., 2013). The mapping was performed with the default settings, and the aligned read counts were generated using the--quantMode function within STAR. The resulting count matrix contained 820880603 uniquely mapped reads, which served as the input for downstream analysis and visualisation, was created by concatenating the assembled reads using Python3 environment.

### Downstream analysis and visualisation

The downstream statistical analysis and plotting were performed in a Python 3.10.12 environment. Gene expression counts were normalised using the pyDESeq2 package (version 0.4.0). Raw counts were transformed using the variance-stabilising transformation (VST) to stabilise variance across samples (Love et al., 2014; Muzellec et al., 2022).

Differential expression analysis was also conducted using pyDESeq2. Genes with an adjusted p-value < 0.1, as determined by the Benjamini-Hochberg method for multiple testing correction, and an absolute log2 fold change > 1 were considered significantly differentially expressed (DEGs). These DEGs were subjected to enrichment analysis to identify overrepresented biological processes using the gseapy package (version 1.0.2).

For the generation of visualisations, the seaborn (version 0.13.2) and pandas (version 2.2.2) libraries in Python were utilised. The correlation matrix was computed using Pearson correlation coefficients on a predefined developmental gene list and visualised with seaborn. The marker gene heatmap was generated using hierarchical clustering with Euclidean distance to group similar expression patterns.

The complete code, including all functions, parameters, and data used for the analysis and visualisation, will be made available on GitHub upon publication.

## Acknowledgements

We thank Sebastian Arnold, Kat Hadjantonakis and Joshua Frenster for cell lines (respectively for the Bra-/-mT, H2B-mCherry;GPI-GFP and H2BemiRFP670 cell lines), Jérôme Gros and Carolina Borja for the 1P-Myosin immunofluorescence protocol, Léo Guignard and Hervé Turlier for discussions. This work was funded by an ERC AdG (MiniEmbryoBlueprint_ 834580) and also a “Maria de Maeztu” Programme for Units of Excellence in R&D (Grant No. CEX2018-000792-M). PC was funded by Spanish Ministry of Science and Innovation and FEDER (grant PGC2018-101251-B-I00). AD was funded by an EMBO postdoctoral fellowship (ALTF 948-2022).

## Competing interests

AMA is an inventor in two patents on Human Polarised Three-dimensional Cellular Aggregates PCT/GB2019/052670 and Polarised Three-dimensional Cellular Aggregates PCT/GB2019/052668.

## Author contributions

AMA and UMF conceived the project, UMF and SB carried out the experimental work and bright field image analysis, PPM performed the bioinformatic analysis, GT and PC carried out the light sheet image analysis, GR and AD made the Nodal and Wnt3 knockout cell lines, and AMA and UMF wrote the paper with contributions from all authors.

---

**Figure S1.**
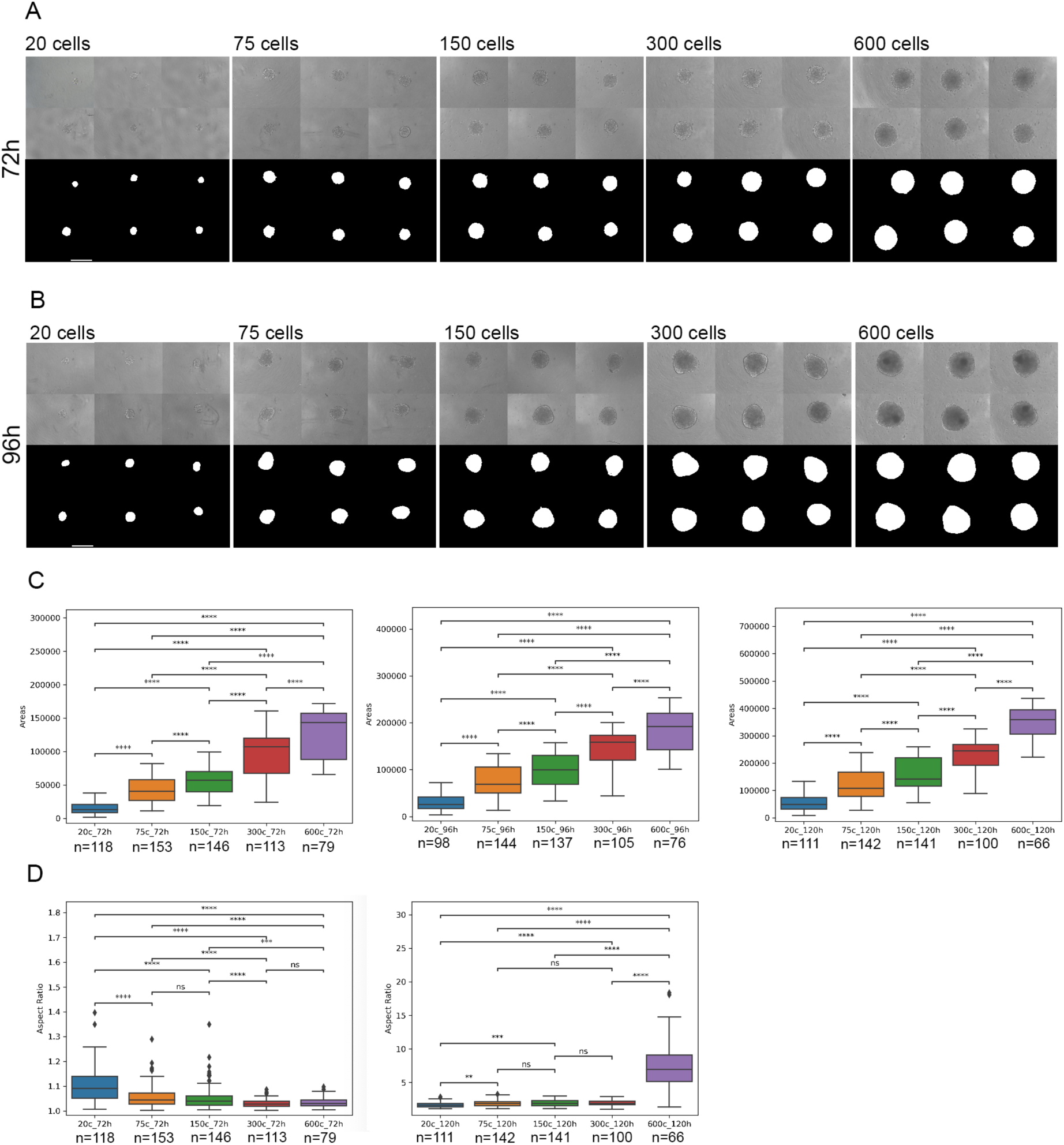
Morphological dynamics of developing E14 gastruloids. (A) E14 derived gastruloids brightfield images and digital masks showcasing the shapes of 72h gastruloids across sizes. (B) E14 gastruloids brightfield images and digital masks showcasing the shapes of 96h gastruloids across sizes. (C) Quantification of gastruloid area (2D segmented digital mask area) across sizes at 72h, 96h and 120h. (D) Quantification of gastruloid aspect ratio (derived from the 2D segmented digital mask) across sizes at 72h and 120h.

**Figure S2.**
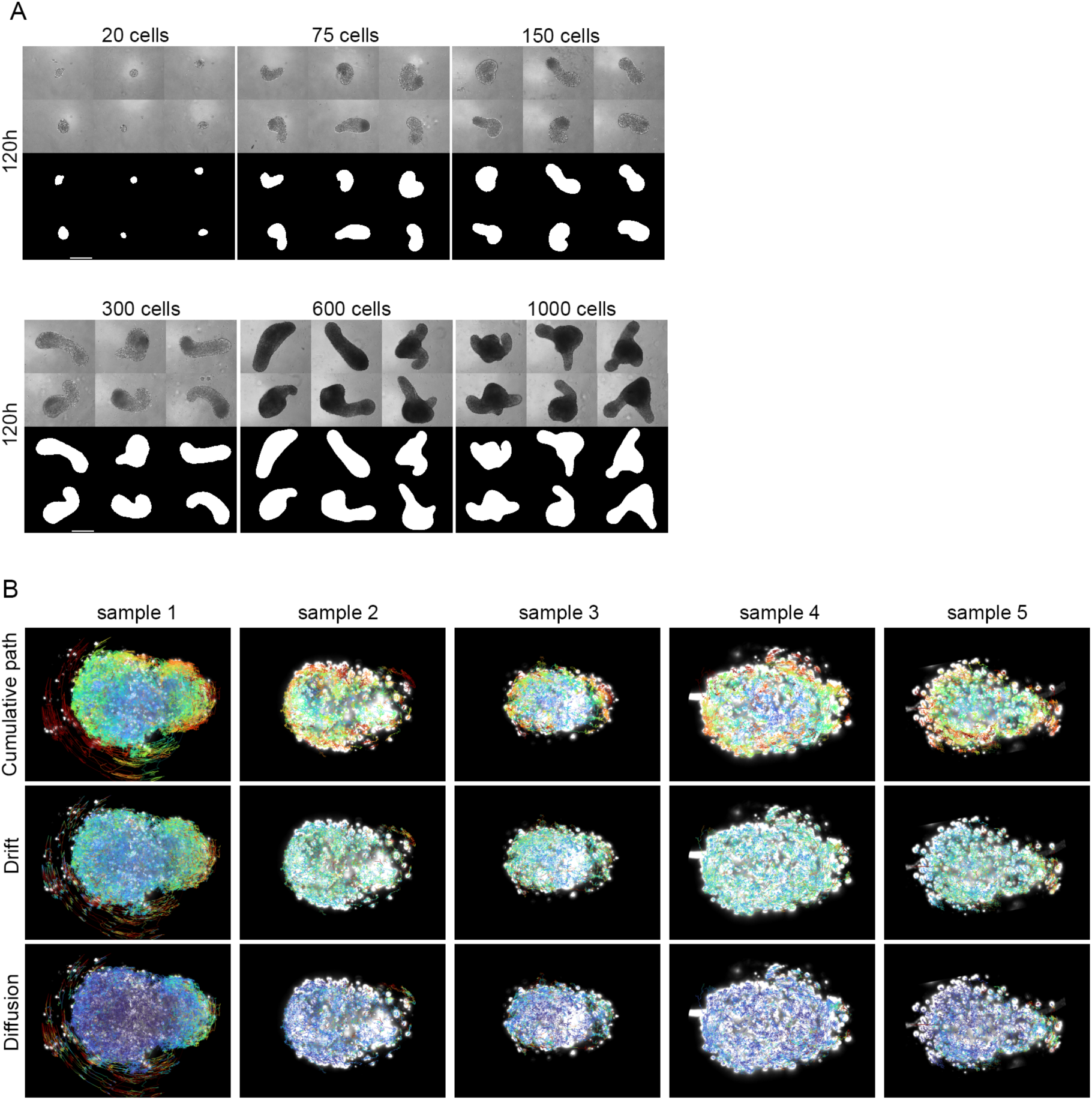
The size range for robust morphological gastruloid development is dependent on the cell line. (A) R1 cells (GPIGFP-H2BmCherry cell line) derived gastruloids brightfield images and digital masks showcasing the shapes of 120h gastruloids across sizes. N=3 replicates. (B) Nuclear movement analysis in 75-cell gastruloids during the early stages of elongation. Data derived from chimeric gastruloids with nuclear sparse-labelling (60% E14 cells + 40% emiRFP670 cells). Tracks colour coded for different motion metrics: path cumulative, drift (directionality modulus) and diffusion (mobility metric). Anterior left and posterior right. Images from 5/6 movies analysed. Red – max values; blue - min values.

**Figure S3.**
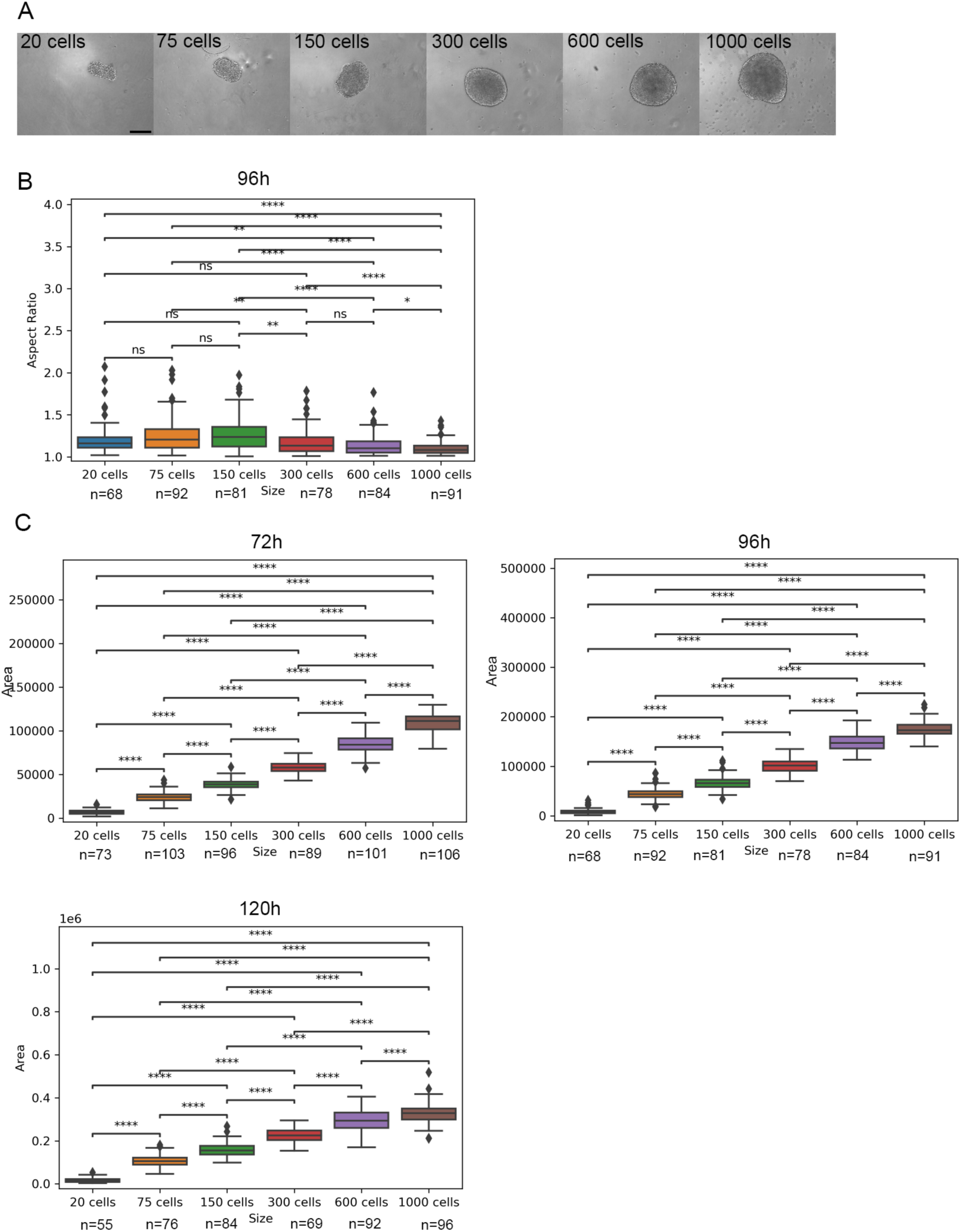
Smaller gastruloids initiate elongation earlier in gastruloids in R1 stem cells. (A) Brightfield images of 96h R1 (GPIGFP-H2BmCherry) gastruloids across sizes. (B) Quantification of aspect ratio of gatruloids across sizes at 96h. (C) Quantification of 2D areas of gastruloids across sizes at 72h, 96h and 120h.

**Figure S4.**
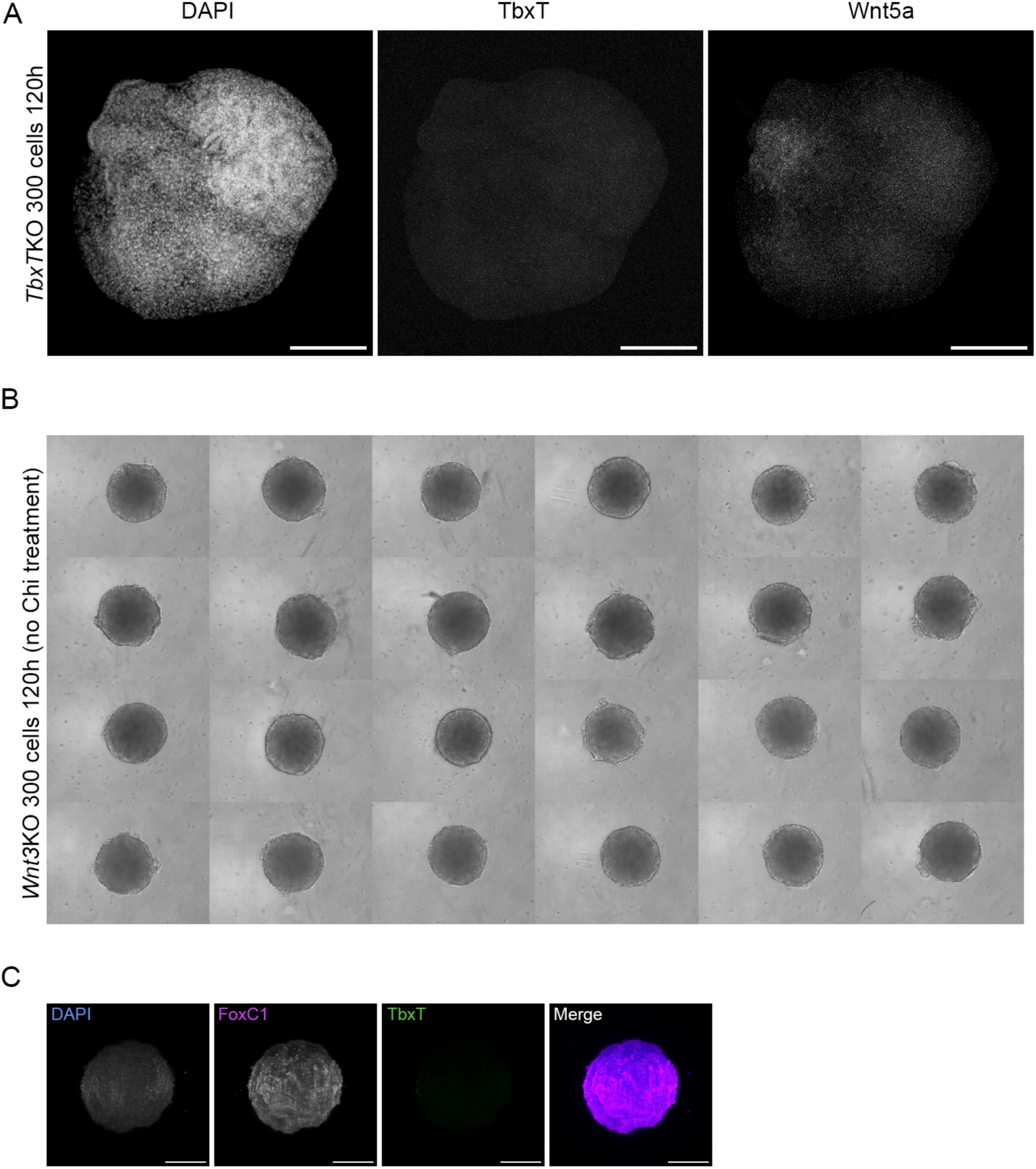
Wnt5a is expressed in 120h *TbxT* mutant gastruloids and Wnt3 is required for TbxT expression. (A) HCR labelling of Tbxt/Bra and Wnt5a in 120h *TbxT*KO gastruloids (control protocol with 3μM Chi treatment between 48-72h). N=3 replicates. (B) Morphological phenotype of *Wnt3*KO aggregates (without 48-72h Chi treatment) at 120h. (C) Patterns of FoxC1 and TbxT expression in *Wnt3*KO aggregates (without 48-72h Chi treatment) at 120h. Scale bar 200μm.

**Figure S5.**
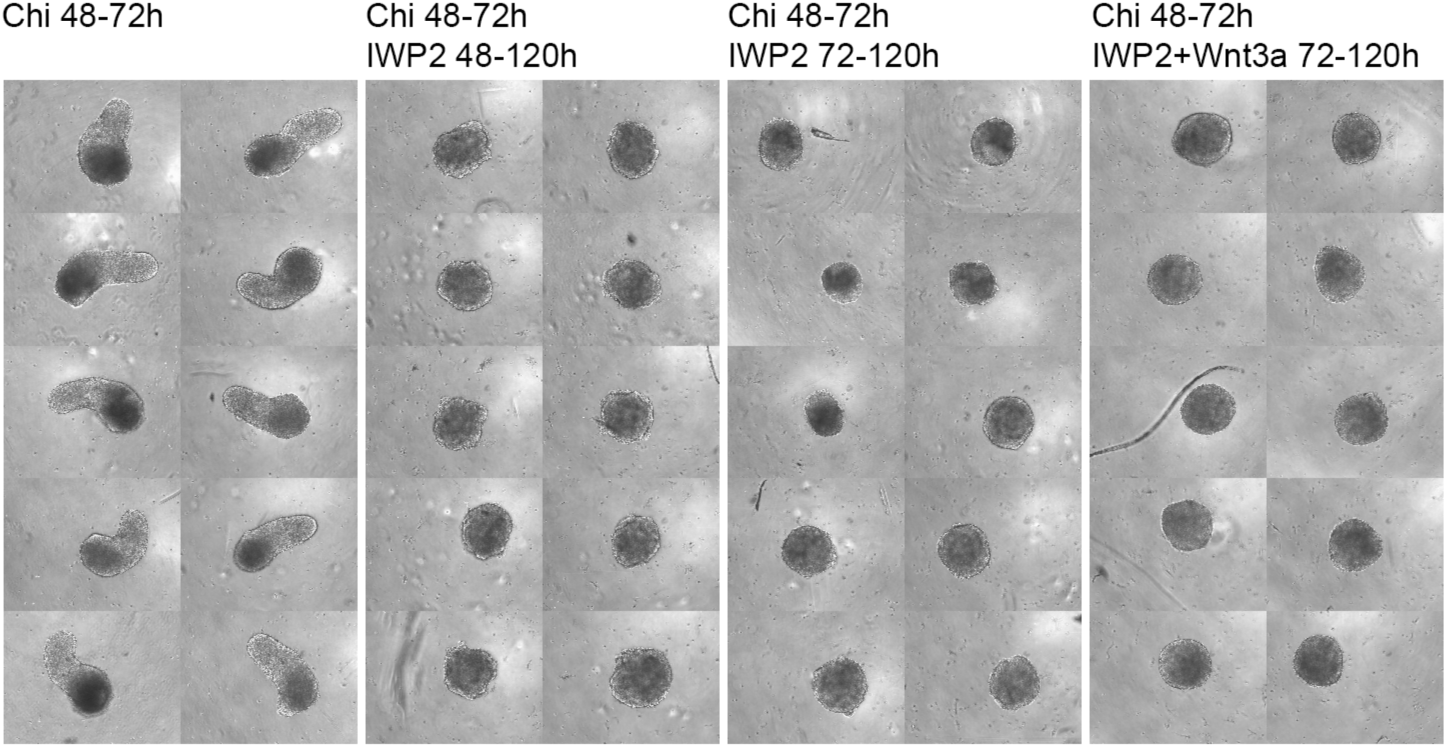
Non canonical Wnt mediated PCP is needed for gastruloid elongation. E14 300-cell 120h gastruloids. Inhibition of Wnt molecules secretion by IWP2 treatment (1μM), prevents elongation in CHIR driven gastruloids. Addition of exogenous Wnt3a (100ng/ml) from 72h does not rescue the IWP2 phenotype of lack of elongation. N=3 replicates.

**Figure S6.**
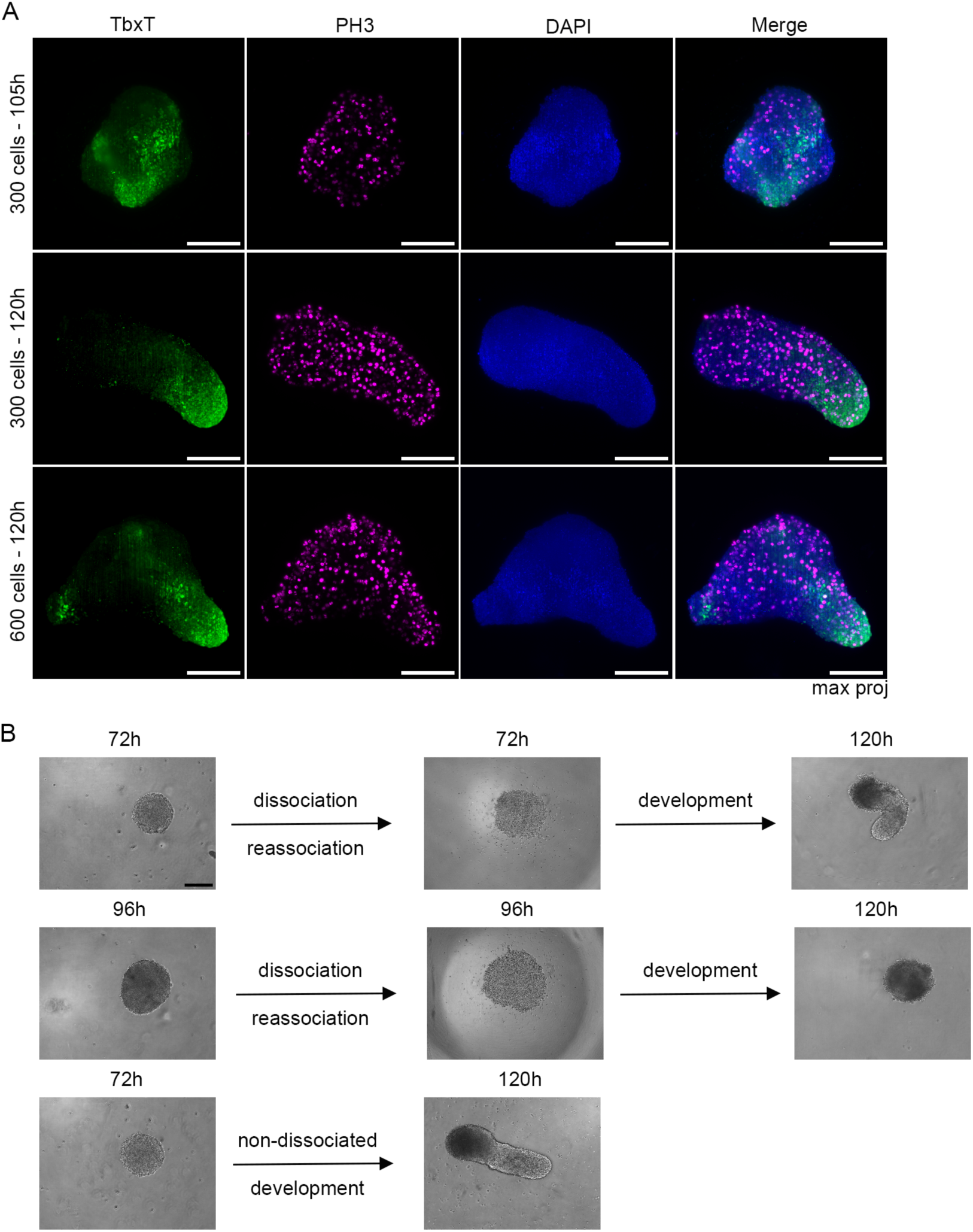
Elongation is not due to anisotropic cell division and the system is relatively adaptable before 96h. (A) Immunostaining of TbxT and Phospho Histone H3 (a marker of mitosis) in 300-cell gastruloids at 105h and in 300-cell and 600-cell gastruloids at 120h. N=3 replicates. (B) Development of gastruloids dissociated and reassociated at 72h, at 96h and non-dissociated. N=3 replicates.

**Figure S7.**
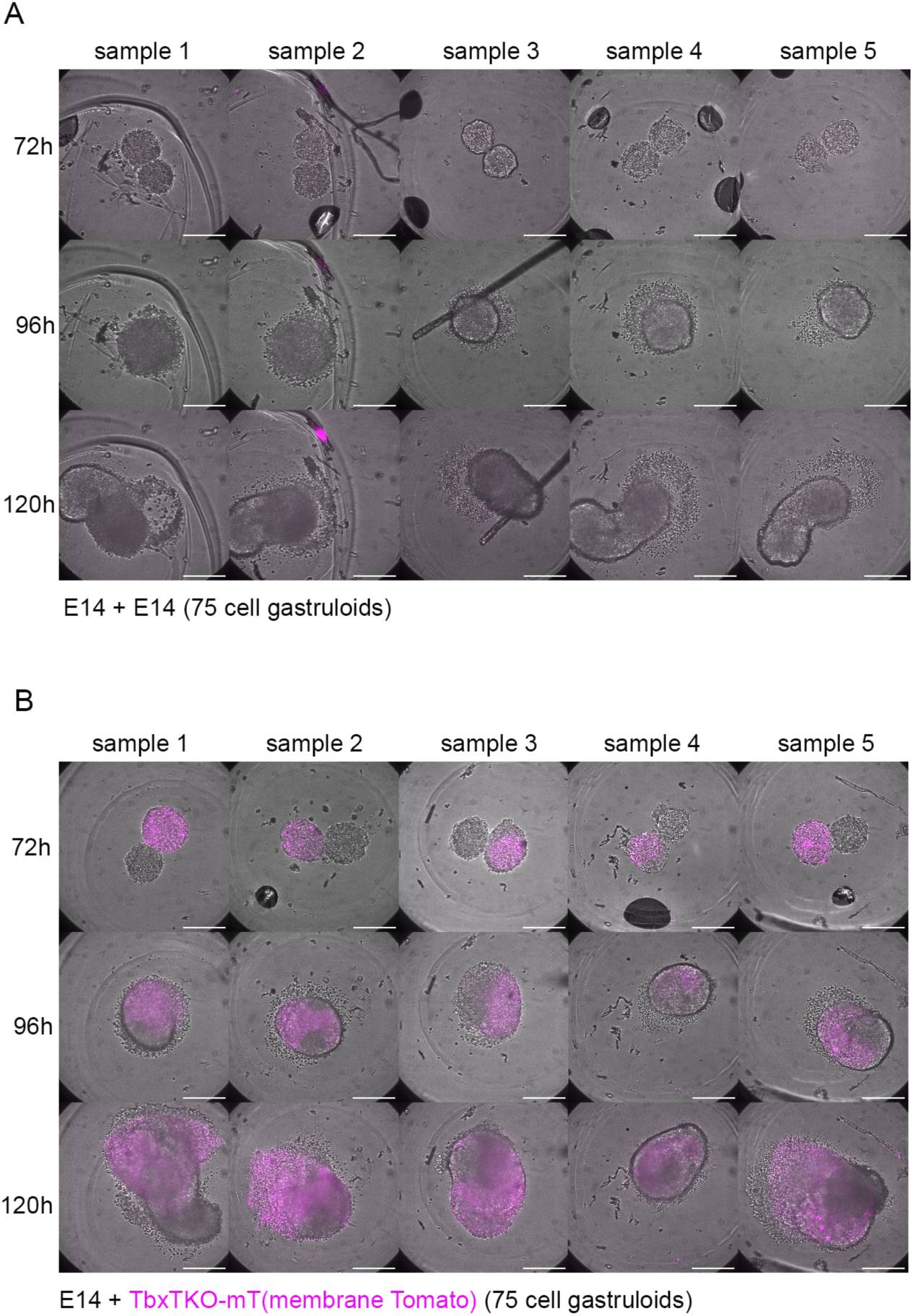
Dynamics of aggregate merging between control and *TbxT* mutant 72h gastruloids. (A) Merging dynamics of E14 75-cell 72h gastruloids. (B) Merging dynamics of E14 with *TbxT* mutant 75-cell 72h gastruloids. N=3 replicates.

**Figure S8.**
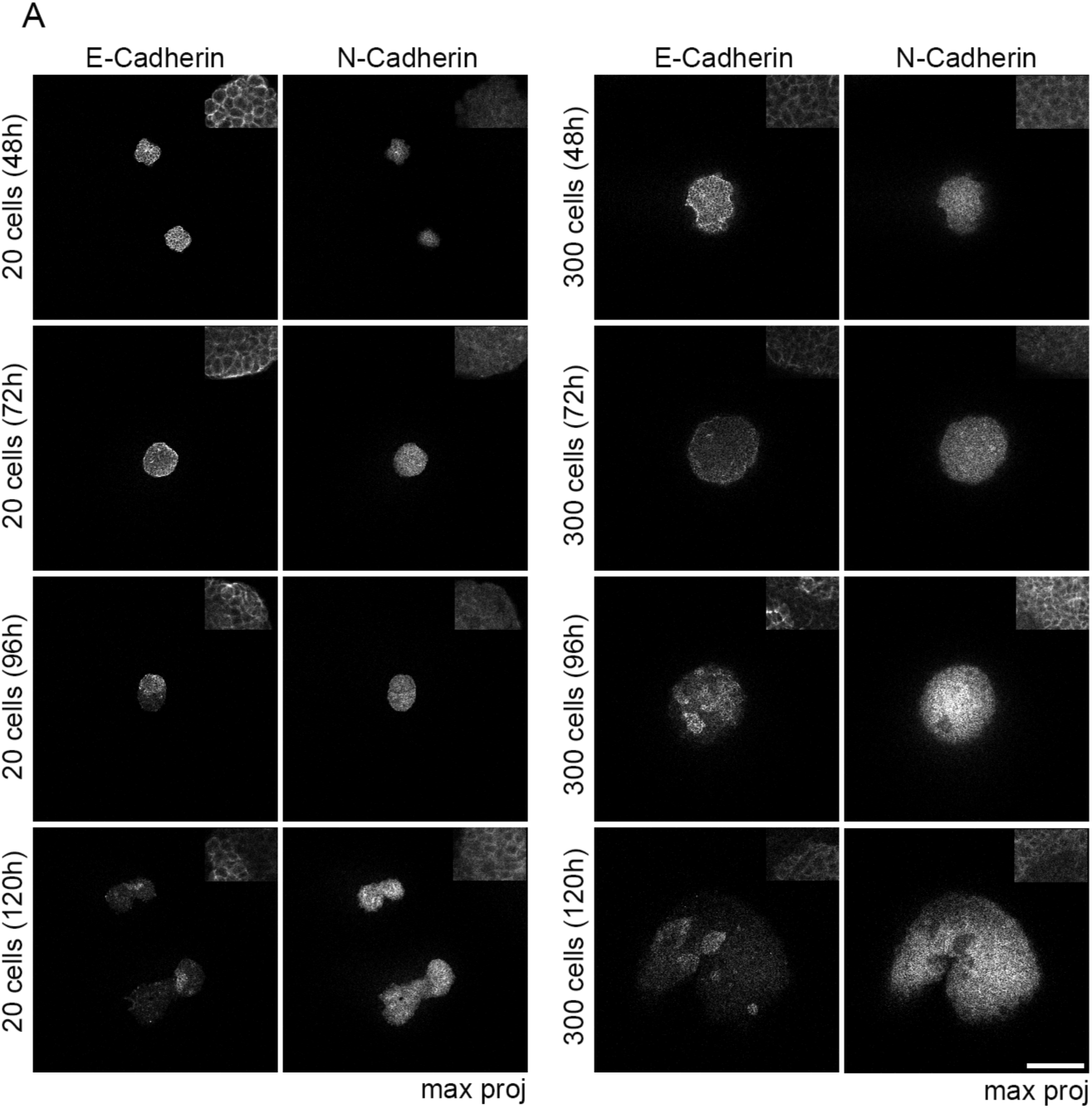
E-Cadherin to N-Cadherin switch during gastruloid development. (A) Evolution of patterns of expression of E-Cadherin and N-Cadherin in 20-cell and 300-cell gastruloids (immunofluorescence). N=3 replicates.

**Figure S9.**
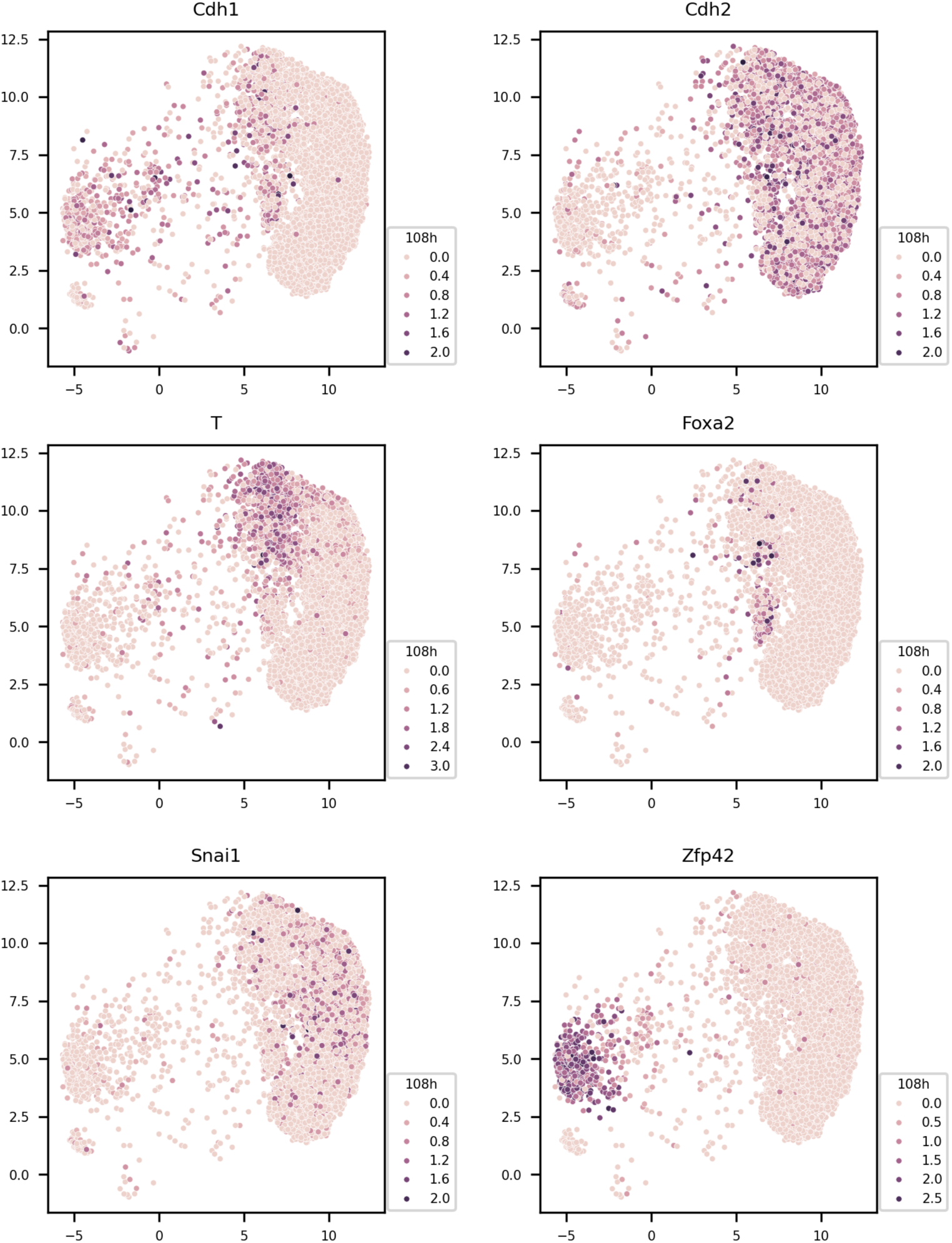
UMAPs of scRNA expression of Cdh1/E-Cadherin, Cdh2/N-Cadherin, T/Bra, FoxA2, Snail and Zfp42 at 108h of gastruloid development. The UMAPs were produced using the publicly released data from the Liberali laboratory (Suppinger et al, 2023).

**Figure S10.**
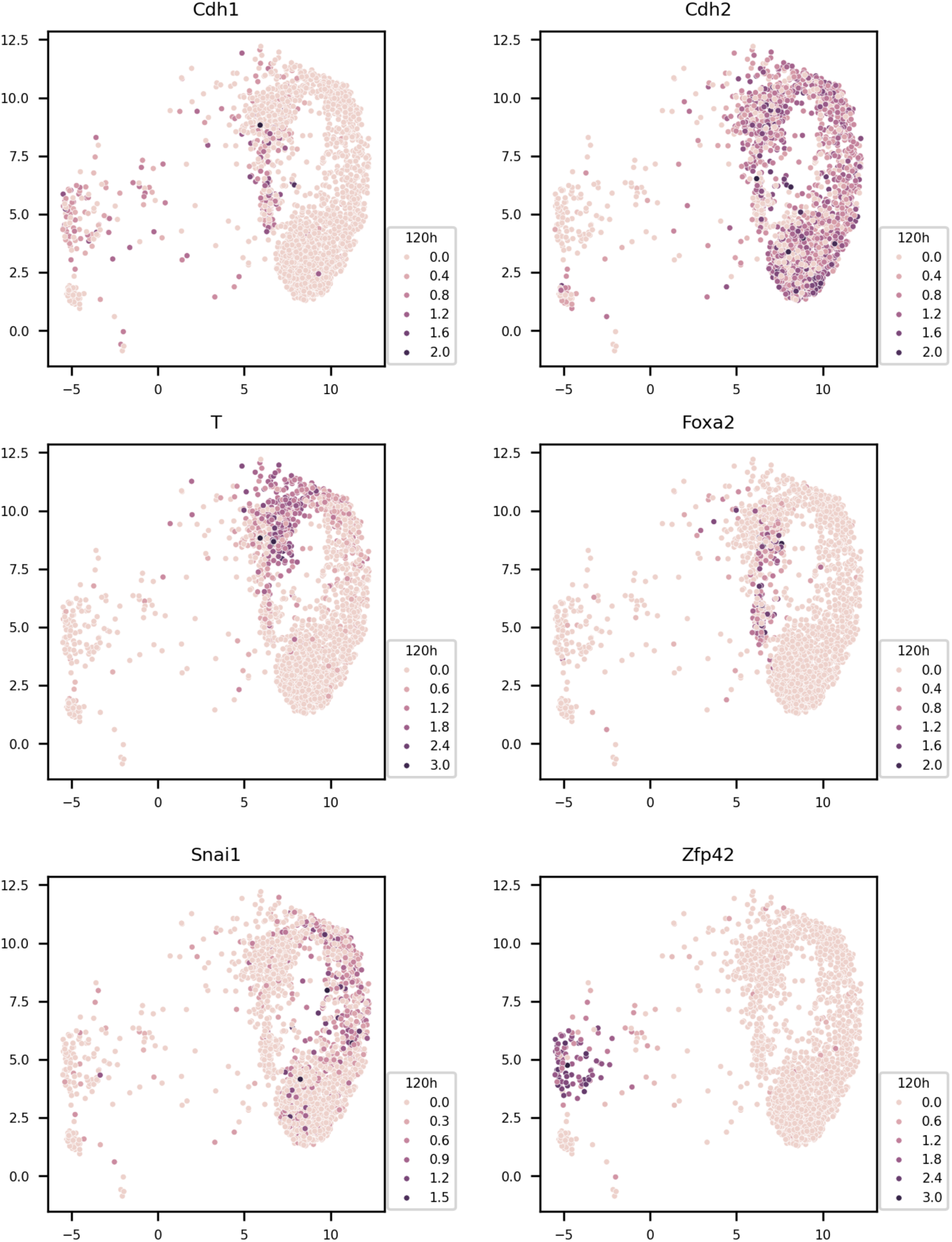
UMAPs of scRNA expression of Cdh1/E-Cadherin, Cdh2/N-Cadherin, T/Bra, FoxA2, Snail and Zfp42 at 120h of gastruloid development. The UMAPs were produced using the publicly released data from the Liberali laboratory (Suppinger et al, 2023).

**Figure S11.**
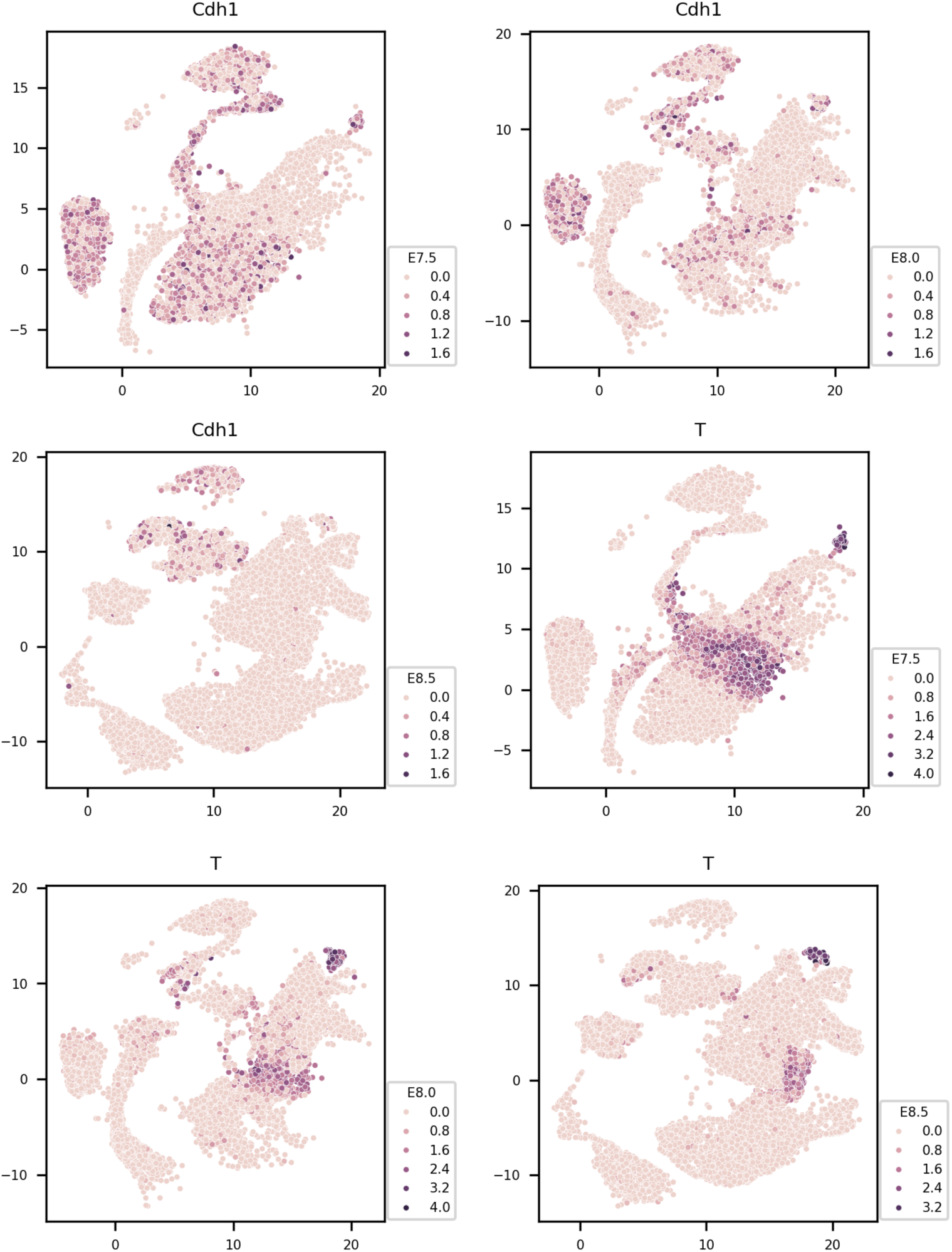
UMAPs of scRNA expression of Cdh1/E-Cadherin and T/Bra at E7.5, E8.0 and E8.5 of mouse embryonic development. The UMAPs were produced using the publicly released data from the Marioni and Göttgens laboratories (Sala et al, 2019).

**Figure S12.**
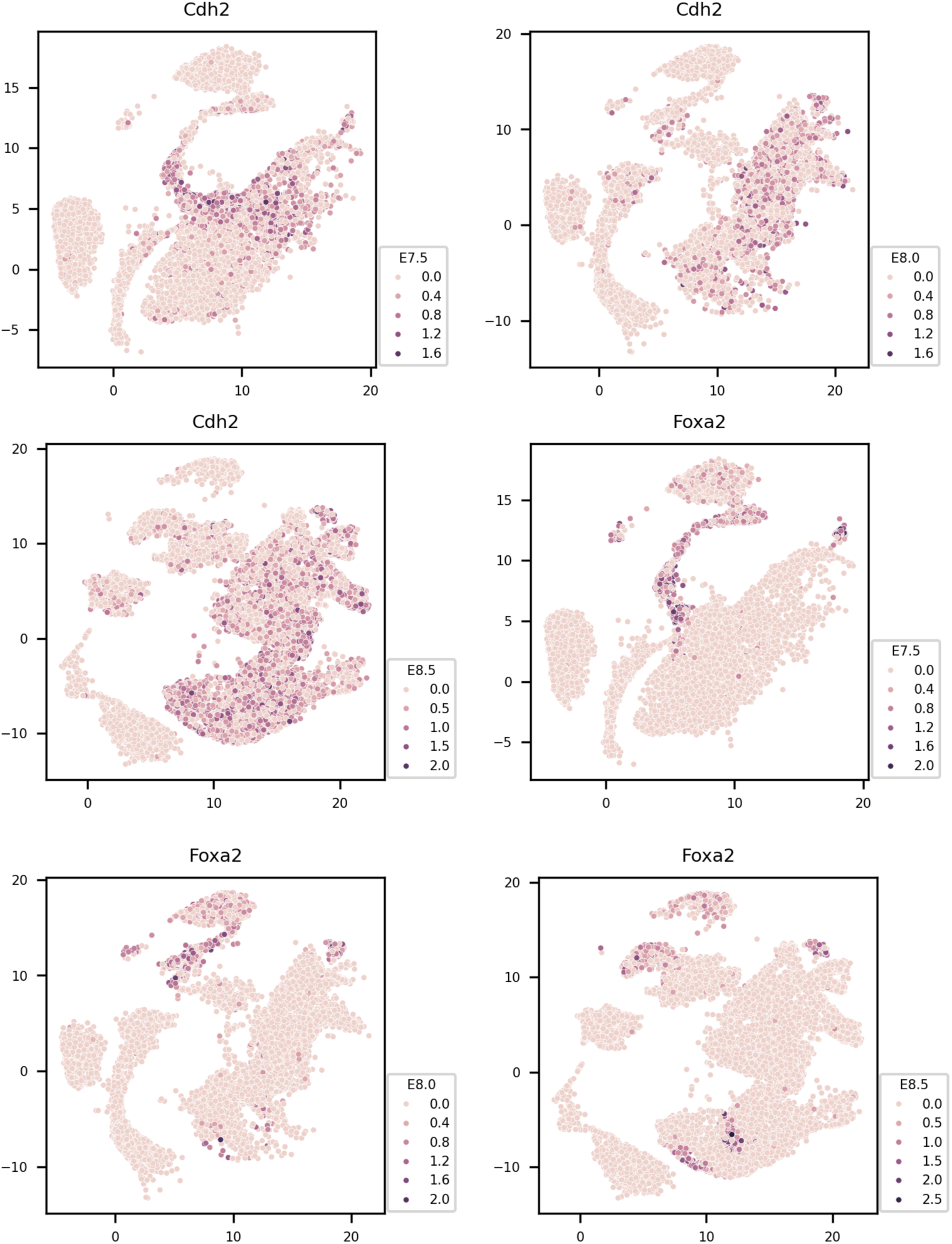
UMAPs of scRNA expression of Cdh2/N-Cadherin and FoxA2 at E7.5, E8.0 and E8.5 of mouse embryonic development. The UMAPs were produced using the publicly released data from the Marioni and Göttgens laboratories (Sala et al, 2019).

**Figure S13.**
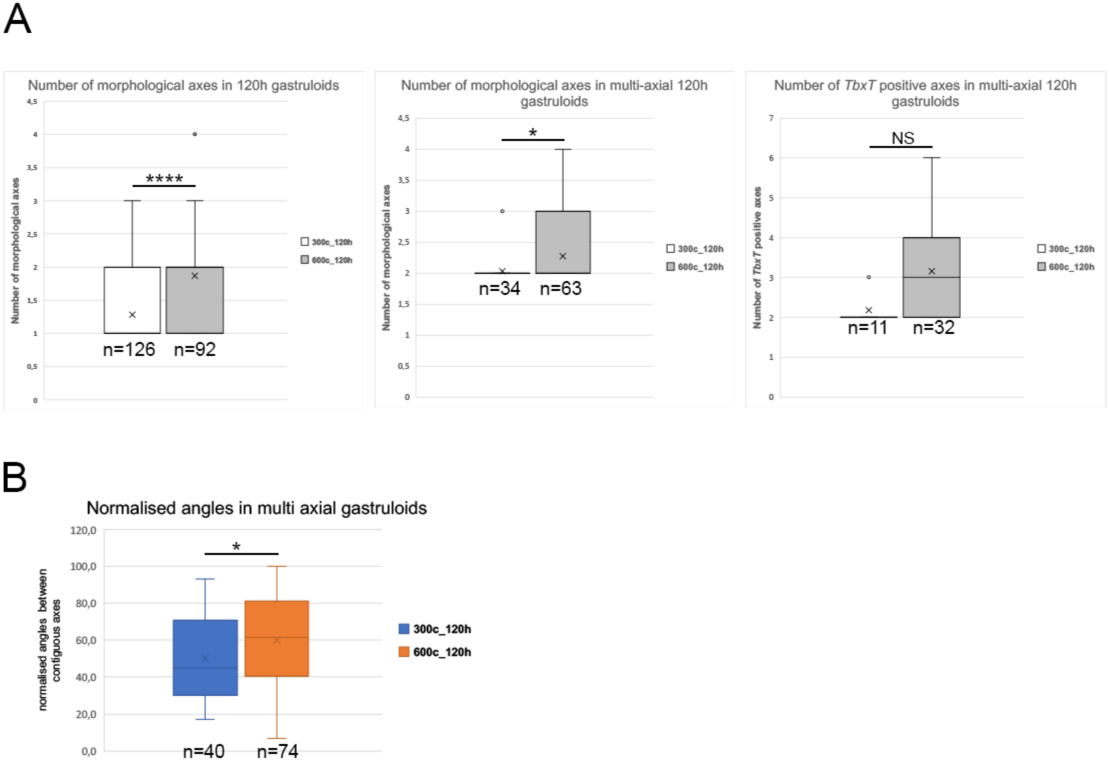
Multi-axis formation in 300 and 600-cell gastruloids. (A) Number of axes formed at 120h in 300-cell and 600-cell gastruloids. (B) Box plot distribution of normalised angles between contiguous axes in 120h 300-cell and 600-cell gastruloids. Only angles smaller than 360° divided by the number of axes were included in the analysis. Normalised angle = (measured angle x 100) / (360 / number of axes).

**Figure S14.**
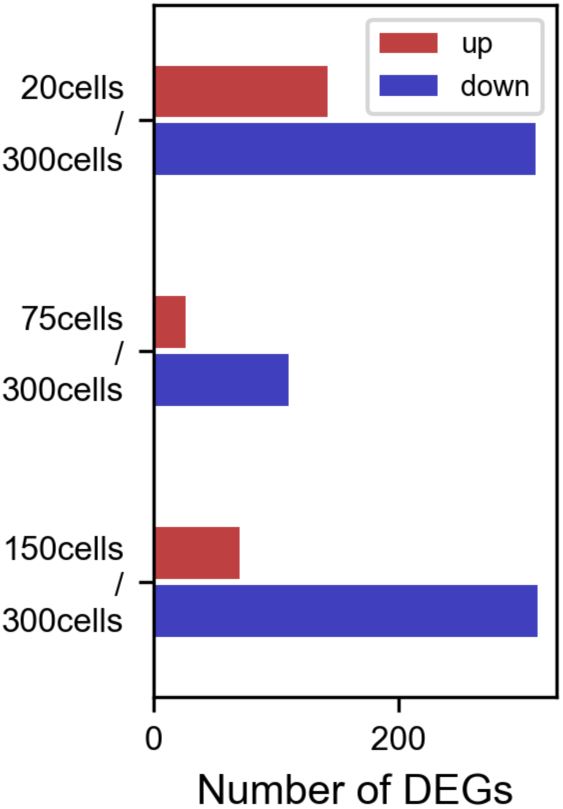
Differential gene expression comparison between 120h gastruloids of different sizes and 120h 300-cell gastruloids. DEG thresholds were set at LFC>1 and p-adjusted value<0.1.

**Figure S15.**
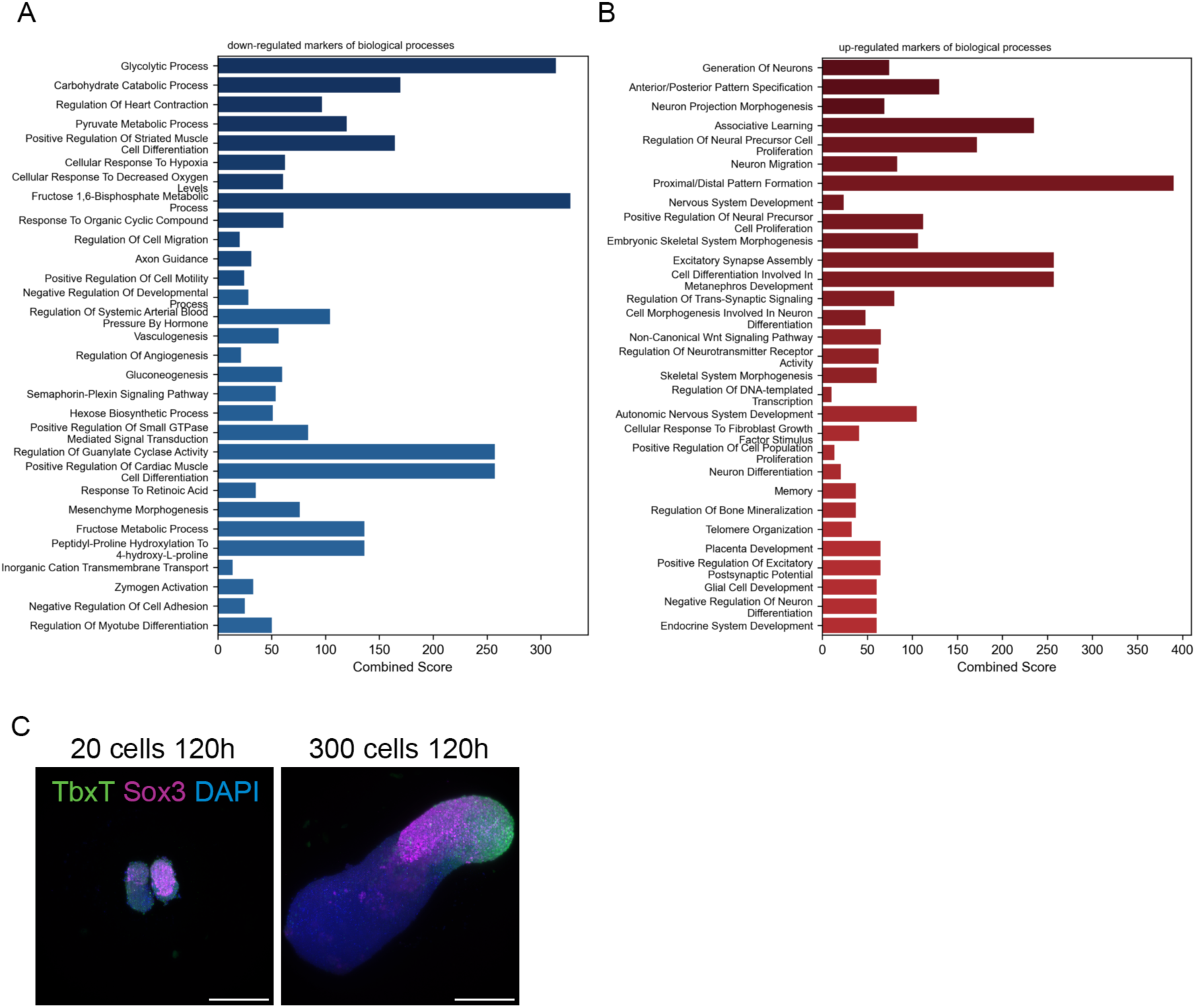
Gene Ontology (GO) analysis of fate and biological processes comparing 20-cell and 300-cell gastruloids at 120h. (A) GO analysis of fate comparing 20-cell and 300-cell gastruloids at 120h. (B) GO analysis of biological processes comparing 20-cell and 300-cell gastruloids at 120h. (C) TbxT and Sox3 expression in 120h 20-cell and 300-cell gastruloids.

**Figure S16.**
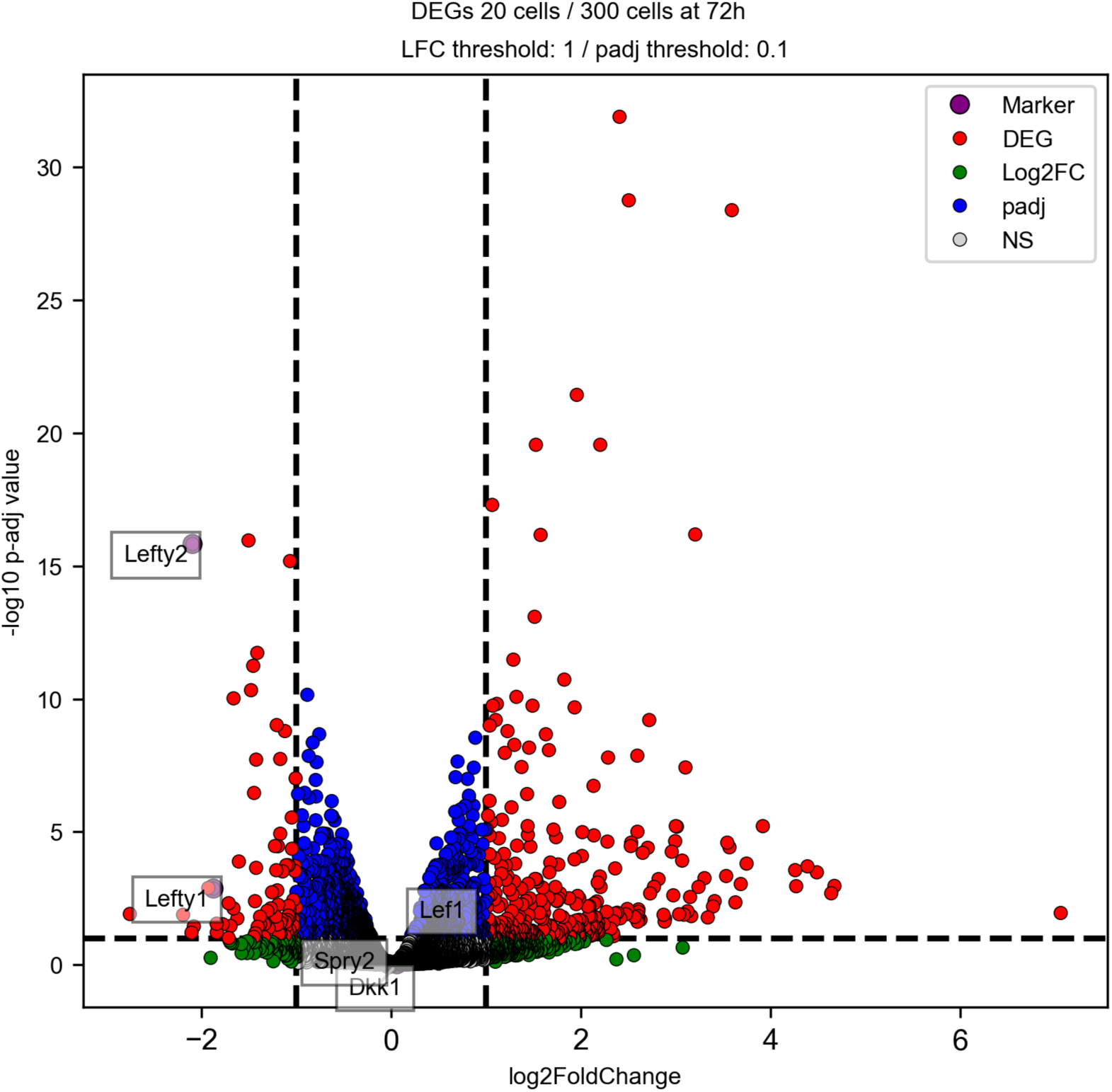
Differential gene expression volcano plot comparing 20-cell and 300-cell, 120h gastruloids highlighting Lefty1, Lefty2, Spry2, Dkk1 and Lef1.

**Figure S17.**
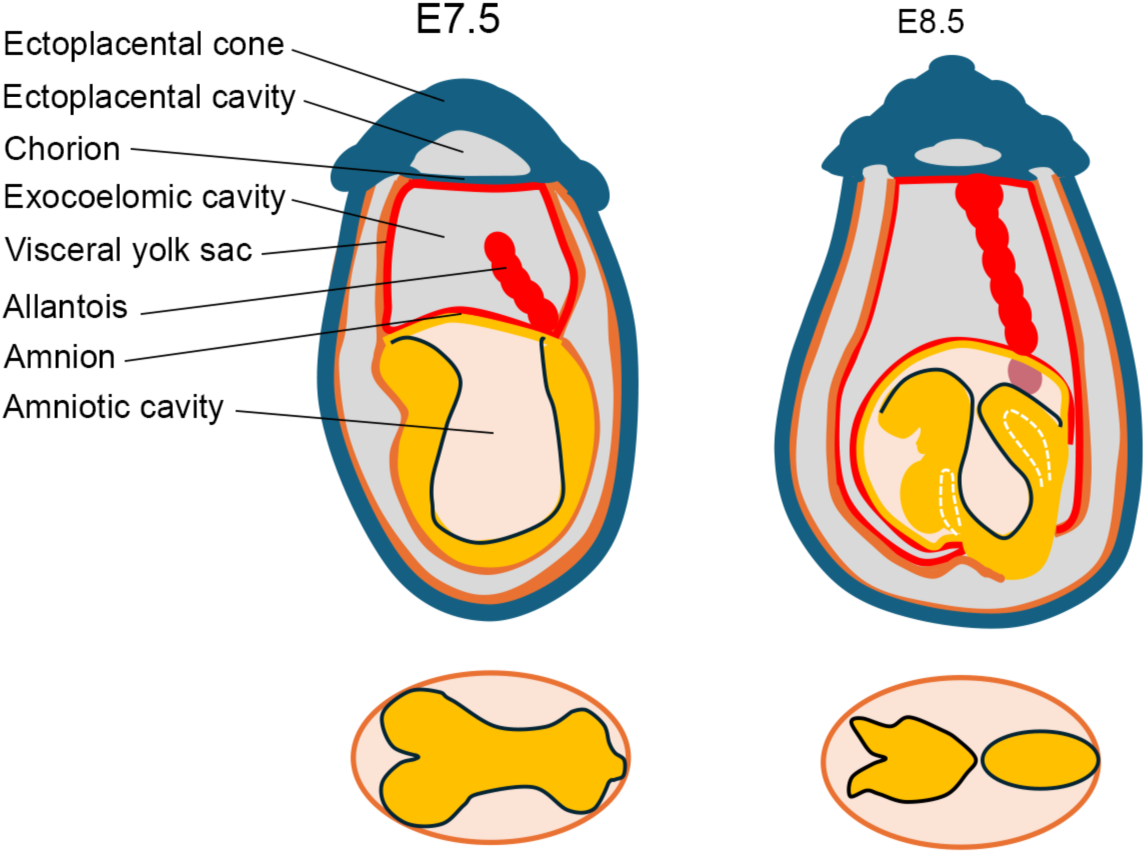
Scheme of the anatomy of a developing mouse embryo.

**Figure S18.**
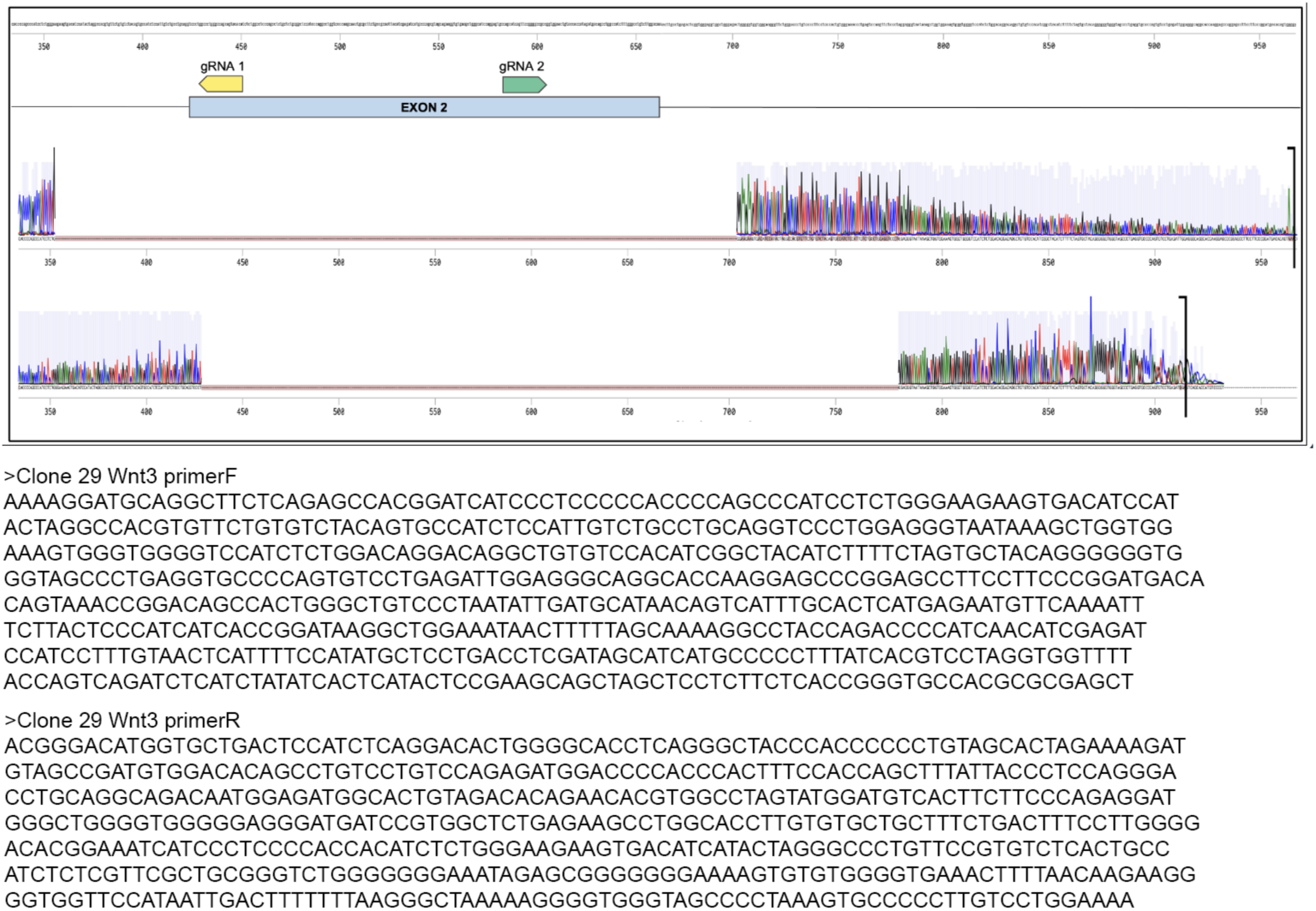
Making of a E14 *Wnt3* mutant line. CRISPR-Cas9 exon2 deletion and frame-shift induction of the *Wnt3* gene.

**Figure S19.**
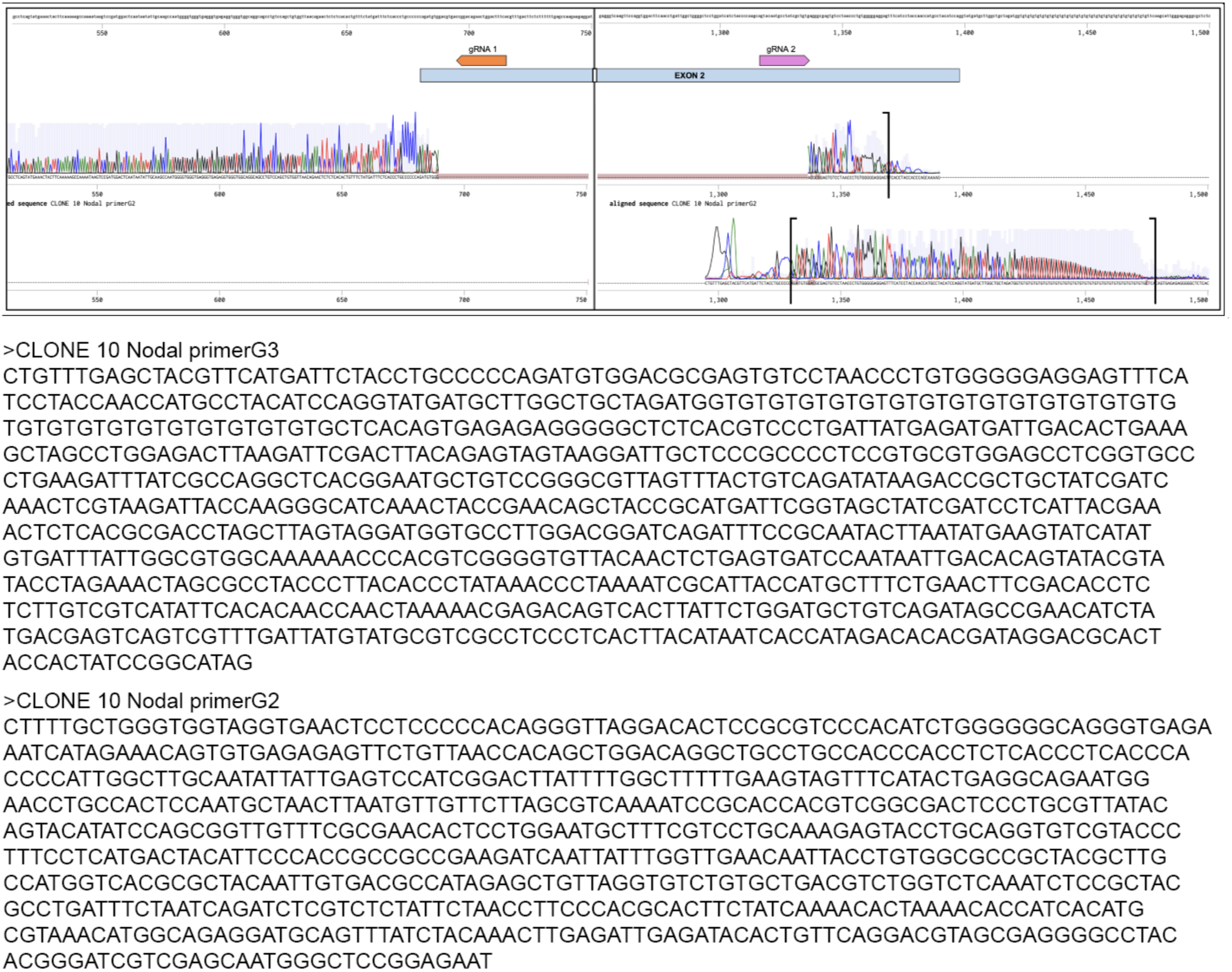
Making of a E14 *Nodal* mutant line. CRISPR-Cas9 exon2 deletion and frame-shift induction of the *Nodal* gene.

**Figure S20.**
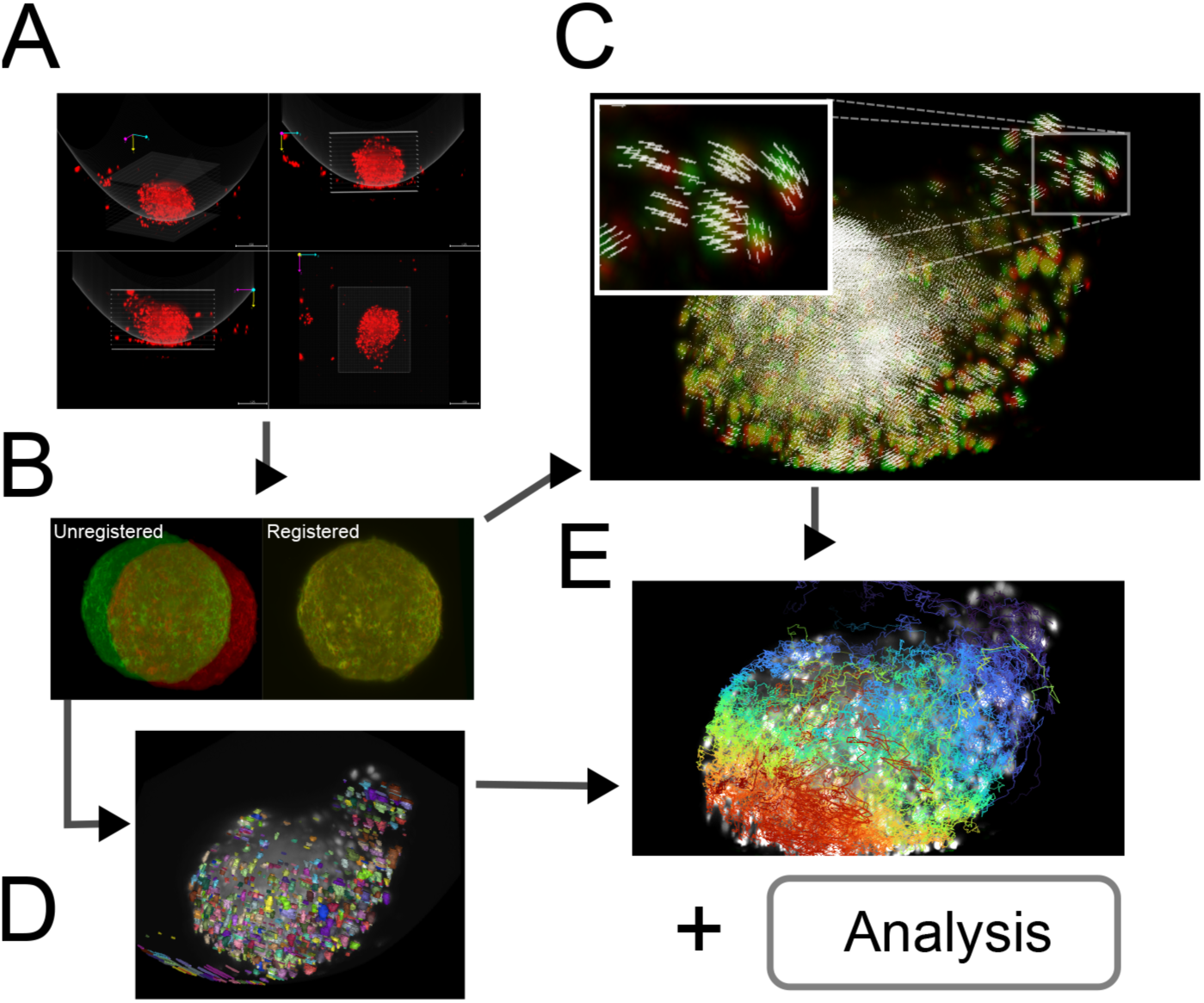
Light sheet data image analysis pipeline. Chimeric E14/E14-emiRFP670 gastruloids (6:4) were imaged on a light sheet microscope with dual illumination and single detection from below (LS1 Viventis). (A) Artifacts like debris and gastruloid extruded cells falling during recording time by gravity, many of which between the sample and the detection objective, can affect registration analysis. To minimise the effect of debris outside the region of the gastruloid, we crop the volume of the gastruloid using a bounding box. To reduce the imapct of extruded cells between the plate and the gastruloid, an empirical parabolic surface is defined, similar to the shape of the sample well, and pixel values this surface are set to zero. (B) Sample movement is corrected by rigid registration. (C) Relative deformations are computed using non-linear registration, which produces a dense vector field. This field is compressed for computational efficiency into a K-NN regressor (See Methods). (D) Cells are segmented from the rigid-registered images using Cellpose. (E) Centroids of the segmented masks are used to backtrack over the non-linear vector field to obtain deformation trajectories of the gastruloid. Over these tracks, different metrics of movement are computed (cumulative path, drift and diffusion, see Methods).

## Movies

**Movie 1.** Bright field time lapse imaging of E14 developing gastruloids with 75 cells (left) and 300 cells (right) starting sizes. Acquired in a Zeiss Axio Observer Z1.

**Movie 2.** Dynamic nuclear motion analysis characterising regional cell movement in a 75 cell gastruloid where multipoles coalesce.

**Movie 3.** Dynamic nuclear motion analysis characterising regional cell movement in a 75 cell gastruloids gastruloids with single pole elongation.

**Movie 4.** SPIM movie of active protrusion dynamics during elongation in 75 cells (left) and 300 cells (right) gastruloids. The cell line imaged is GPIGFP;H2BmCherry. Imaged on a Viventis LS1.

**Movie 5.** Magnified SPIM movie of active protrusion dynamics during elongation in 75 cells gastruloids. The cell line imaged is GPIGFP;H2BmCherry. Imaged on a Viventis LS1.

**Movie 6.** Bright field time lapse imaging of E14 control (left) and Cytochalasin D treated (right) developing gastruloids. Cytochalasin treatment of 0.05 ng/ml from 90h after cell seeding. Acquired in a Zeiss Axio Observer Z1.

**Movie 7.** Time lapse imaging of 20 cells merging aggregates, labelled and not labelled with Hoechst, from 72h. Imaged on a Viventis LS1.

**Movie 8.** Time lapse imaging of 75 cells merging aggregates, labelled and not labelled with Hoechst, from 72h. Imaged on a Viventis LS1.

**Movie 9.** Transmitted light time lapse imaging of morphological dynamics during elongation. Imaged on a Viventis LS1. Chimeric E14 WT (60%) and E14 H2BemiRFP670 (40%) gastruloids.

## Notes

### Summary of Updates

Competing interests statement and authors list.

